# Peripheral lysosomes recruit PLEKHG3 to focal adhesions and restrain protrusion dynamics

**DOI:** 10.1101/2023.02.01.526449

**Authors:** Rainer Ettelt, Ana-Maria Sandru, Georg Vucak, Sebastian Didusch, Biljana Riemelmoser, Karin Ehrenreiter, Markus Hartl, Lukas A. Huber, Manuela Baccarini

## Abstract

Lysosomes are dynamic organelles regulating metabolic signaling by recruiting cytosolic molecules to protein platforms on their limiting membrane. We used proximity labeling to define interactors and vicinal proteins of LAMTOR3, a component of the Ragulator scaffold that controls mTORC1 signaling and lysosome positioning. The screen has yielded several previously unappreciated interactors, including an actin remodeling network. Here, we characterize the RhoGEF PLEKHG3 as a LAMTOR3 vicinal protein colocalizing with peripheral lysosomes and cortical F-actin at focal adhesion sites. Forced peripheral dispersion of lysosomes drives PLEKHG3 accumulation at focal adhesions and decreases protrusive activity in both wild-type and PLEKHG3-deficient cells. Thus, lysosome positioning governs both PLEKHG3 localization and protrusive activity, yet the protrusion changes can occur independently of PLEKHG3.

## Materials Methods

### Cell culture

Human embryonic kidney (HEK) 293T, Flp-In™ T-REx™ 293 (Invitrogen™) and Human HeLa cervical carcinoma (HeLa) cells were cultured in DMEM (D5796; Sigma-Aldrich) supplemented with 10% FBS, and 100 U/ml penicillin–streptomycin at 37°C, 5% CO_2_. Stable GFP and GFP-PLEKHG3 HeLa cells were cultured under similar conditions with 100 µg/ml hygromycin B (H3274; Sigma-Aldrich).

### Transfection

Transfections were performed with 1 µg DNA in 6-well plates and 400 ng DNA in 24-well plates. Cells were starved in serum-free medium for 30 min prior to transfection. DNA and PEI transfection reagent were incubated and mixed in plain DMEM (D5796; Sigma-Aldrich). Starvation medium was exchanged with 9 parts full medium and 1 part transfection mix. Cells were incubated for about 24 hours at culture conditions. For live imaging constructs, transfections were performed with 300 ng mTagBFP2-BFP-LifeAct-7 (#54602; Addgene) and 500 ng mCherry/mCherry-KIF1A DNA in 8-well glass bottom plates (IBIDI) using the Lipofectamine 3000 transfection kit (L3000150; Invitrogen) according to the manufacturer’s instructions. The transfection mix was added in full medium, which was replaced the next day by imaging medium (see Live Cell imaging section).

### siRNA-based knockdown

Oligo duplex siRNA against human PLEKHG3 (SR308671) were purchased from OriGene. Final siRNA concentrations of 20, 40 or 60 nM were used for silencing (indicated in figures) and incubated for 36-48 hours. Transfections were performed using Lipofectamine™ RNAiMAX transfection reagent (Invitrogen; 13778-150) according to manufacturer’s instructions.

### Generation of a PLEKHG3 KO cell line

Using CRISPR/Cas9, we evaluated 10 sgRNAs for efficient PLEKHG3 KO (see Table 1) based on the output of the sgRNA tool VBC Score [1].

**Table 1:**
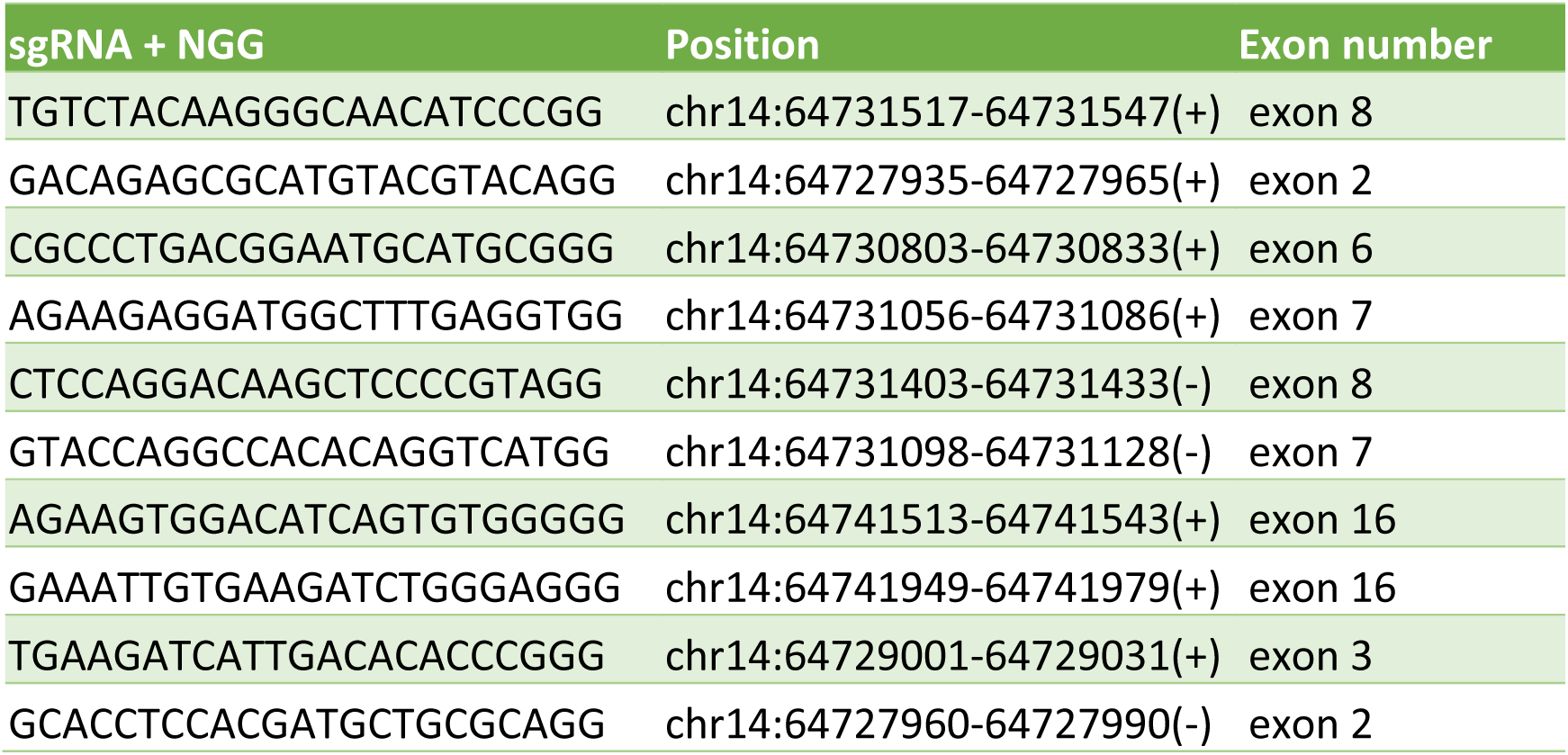
PLEKHG3 CRISPR/Cas9 KO guides. Top 10 PLEKHG3 guides from VBC Score tool with sequence, chromosome and exon position given.

sgRNAs were subcloned into the pX458 backbone (Addgene #48138) and applied according to manufacturer’s instructions. Cells were transfected with the indicated sgRNAs and FACS bulk sorted 24 hours post transfection using a BD FACSAria IIIu Cell Sorter. Cells positively transfected expressed a green fluorescent protein (GFP) in the cytoplasm based on which a FITC sort was applied, selecting the cells with high expression levels. The sorted positive cells were carefully centrifuged, resuspended in growth medium and seeded into 24 or 6 well plates based on the number of sorted cells. The transfection and FACS sorting steps were repeated after a few days, to ensure a high efficiency of the knock-out in the clones. After successful growth of 7 out of 10 guides, cells were seeded for Immunoblot analysis and KO efficiency was tested based on intensity reduction of PLEKHG3 band (Figure S2G). Successful KO cells were kept in culture and used for subsequent experiments.

### Generation of DNA constructs and stable cell lines

Genes of interest for transient transfection were cloned into a pCDNA3.1 vector by Gibson assembly. In brief, pCDNA3.1 vector was linearized by restriction digested with BamHI and EcoRI. mCherry insert was amplified by PCR from pcDNA5-MTS-TagBFP-P2AT2A-EGFP-NLS-P2AT2A-mCherry-PTS1 (87829; Addgene) using the primers 5’-caagcttggtaccgagctcggatccATGGTGAGCAAGGGCGAG-3’ (forward) and 5’-agtgtgatggatatctgcagaattcttaCTTGTACAGCTCGTCCATGC-3’ (reverse); mCherry-KIF1A insert was amplified from pCW57.1-mCherry-KIF1A (inhouse, built from eGFP-KIF1A (172206; Addgene)) using the primers 5’-caagcttggtaccgagctcggatccgccaccATGGTGAGCAAGGGCGAG-3’ (forward) and 5’-agtgtgatggatatctgcagaattcTCAGACCCGCATCTGCGC-3’ (reverse); RUFY3-mCherry insert was PCR amplified from RUFY3 cDNA with primers 5’-caagcttggtaccgagctcggatccGCCACCATGTCTGCTCTG-3’ (forward) and 5’-tgctcaccatTCCTCCTGATGGGCTGGTAG-3’ (reverse) and mCherry from pcDNA5-MTS-TagBFP-P2AT2A-EGFP-NLS-P2AT2A-mCherry-PTS1 (87829; Addgene) using the primers 5’-atcaggaggaATGGTGAGCAAGGGCGAG-3’ (forward) and 5’-agtgtgatggatatctgcagaattcTTACTTGTACAGCTCGTCCATG-3’ (reverse). All inserts were fused into pCDNA3.1 backbone by Gibson assembly according to the manufacturer’s instructions. pEGFPC1-KIF1B was constructed by Gibson assembly of PCR amplified human KIF1B derived from cDNA using the primers 5’-gatctcgagctcaagcttcgaattctATGTCGGGAGCCTCAGTG-3’ (forward) and 5’-tcagttatctagatccggtggatccTTAGTATTTCGACTGGCTCGG-3’ (reverse) with BamHI and EcoRI digested pEGFPC1 backbone. GFP and GFP-PLEKHG3 plasmids were constructed by Gibson assembly of BamHI and EcoRI restriction digested pCSII-hygro backbone with GFP using the primers 5’-acacgctaccggtctcgacgaattcGCCACCATGGTGAGCAAG-3’ (forward) and 5’-ctagcttaagttagttaacggatccTCACTTGTACAGCTCGTCC-3’ (reverse) or GFP-PLEKHG3 from pEGFP-GFP-PLEKHG (inhouse) using the primers 5’-acacgctaccggtctcgacgaattcGCCACCATGGTGAGCAAG-3’ (forward) and 5’-ctagcttaagttagttaacggatccTCAACCGACAGAGTTCAAGG-3’ (reverse). Stable HeLa cells expressing GFP or GFP-PLEKHG3 were generated by lentiviral transduction and maintained in hygromycin B selection (100 µg/mL). L3 KO cell lines were generated using the CRISPR/Cas9 protocol [2]. In brief, three L3 guides were designed with the benchling CRISPR online tool two of which located in exon 4 (ATCTCTATCTGACACAACAA; TATCTGACACAACAATGGCA) and one in exon 5 (GCAAAAGTGGATAAGAAACC). Single guide RNAs were cloned into pSpCas9(BB)-2A-GFP (PX458) (48138; Addgene) and transfected into HeLa cells. 24 hours post transfection cells were sorted via FACS (BD FACSMelody Cell Sorter) for FITC-positive cells. Bulk sorts were expanded and analysed via Immunoblot for successful L3 deletion. A second round of transfection was necessary to achieve a 95-99% reduction in L3 expression.

### Immunofluorescence

Immunofluorescence was performed as follows: Cells were fixed in 4% formaldehyde in PBS for 10 min, washed three times with 1x PBS and permeabilized with 0.02% Saponin + 3% BSA (w/v) in PBS for 20 min at room temperature. The following primary antibodies diluted in permeabilization solution were used for incubation overnight at 4 °C: rabbit anti-V5 (1:500; #13202), rabbit anti-LAMP1 (1:500; #9091; used in Figures 3F, 4A-B, S3E), rabbit anti-LAMTOR4 (1:500; #13140) were purchased from Cell Signaling Technology; rabbit anti-V5 tag (1:500; V8137) and mouse anti-Vinculin (1:400; V9131) from Sigma-Aldrich; additional antibodies were rabbit anti-PLEKHG3 (1:200; PA5-66764; Invitrogen), mouse anti-paxillin (1:200; 610055; BD Biosciences, discontinued project; used in Figures 4A-B, S3H), rabbit anti-Paxillin (1:1000; ab246718; Abcam; used in Figure 5F) and mouse anti-LAMP1 (1:500; sc-20011; Santa Cruz; used in Figures 3B-D, 5A + F, S1A + F, S2E + H, S5D + F + I). Cells were washed three times with 1x PBS and incubated with the appropriate secondary antibodies conjugated with Alexa Fluor 488, 594, 647, 568 (1:1,000; Thermo-Fisher Scientific) and Alexa Fluor 405 (1:250; Thermo-Fisher Scientific) for 1 h at room temperature. After incubation with DAPI for 10 min (in some cases without DAPI) coverslips were mounted on glass slides with Prolong Gold antifade reagent (P36934; Thermo-Fisher Scientific). Confocal images were captured on an LSM 700 microscope (Zeiss) using a 40x Plan-Apo/1.3 NA oil objective at room temperature with a PMT detector. The acquisition software was Zeiss ZEN 2012 SP5. For medium throughput imaging (indicated in figure legends) slides were imaged on a Slide Scanner VS120 (Olympus) with a 40x dry objective and a CCD camera via the software OlyVIA. High content screening was performed at TIGEM (Naples, Italy) on an Operetta (PerkinElmer) equipped with a Spinning Disk confocal setup using a 20x/0.4 NA air objective and a sCMOS camera at room temperature. All images were analyzed in ImageJ by determining regions of interest (ROIs) in the respective channels. Images are presented with adjusted brightness and contrast. The same adjustments were made in ImageJ for every experiment.

**Figure 1:**
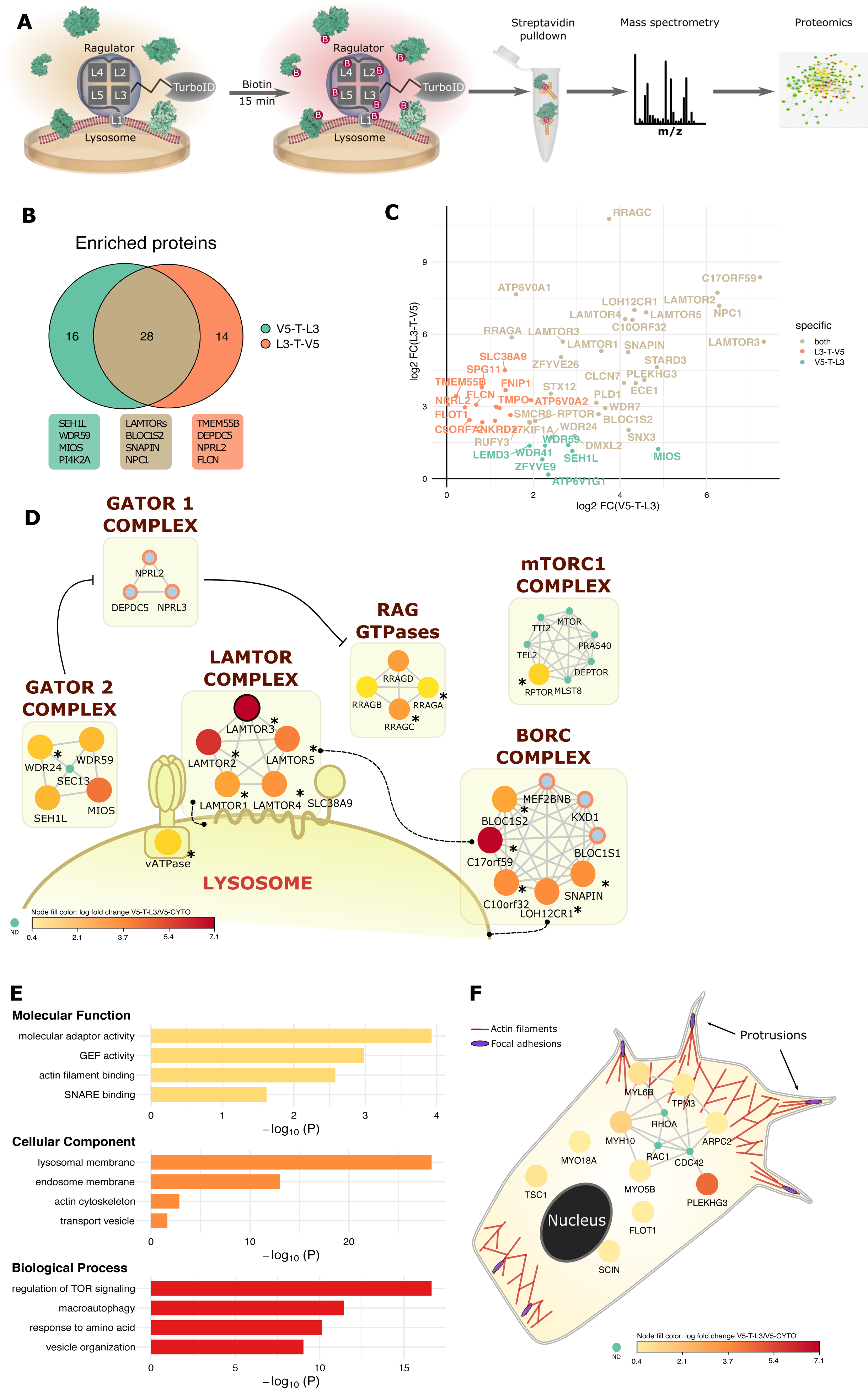
Bioinformatics analysis of L3-TurboID PDL screen. **A)** Workflow of TurboID screens. Cells were treated with biotin for 10-15 minutes prior to lysis and isolation of biotinylated proteins and mass spectrometry. **B)** Venn-diagram of complementary TurboID screens using C-terminally (L3-T-V5) and N-terminally (V5-T-L3) tagged L3. Shown are hits significantly enriched in L3 over the respective cytosolic controls and in V5-LYSO over V5-CYTO (log2 FC ≥ 1.49 and adj. p-value ≤ 0.05). **C)** Fold change plot combining the results of both screens. **D)** Network representation of the complementary L3-TurboID datasets. Depicted proteins are regulators of mTORC1 signaling and well-described interactors of the Ragulator complex. For the hits detected by V5-T-L3 (outlined in black), color coding indicates fold change over V5-CYTO. Proteins detected only by L3-T-V5 are shown as light blue nodes circled in orange. Asterisks mark proteins detected in both screens. Proteins that are part of the networks identified in our screen but not detected by either bait are shown as green dots. Broken lines indicate protein-protein or protein-membrane interactions. **E)** Gene ontology overrepresentation analysis (ORA) of proteins significantly enriched in V5-T-L3 over V5-CYTO. GEF, guanine nucleotide exchange factor. **F)** network and functional representation of V5-T-L3 hits enriched in the GO-terms actin filament binding and actin cytoskeleton, plus manually curated hits. Color coding indicates enrichment over V5-CYTO-TurboID. Actin filaments are shown in red and FAs in purple. Proteins not detected by V5-T-L3 are shown as green dots.

**Figure 2:**
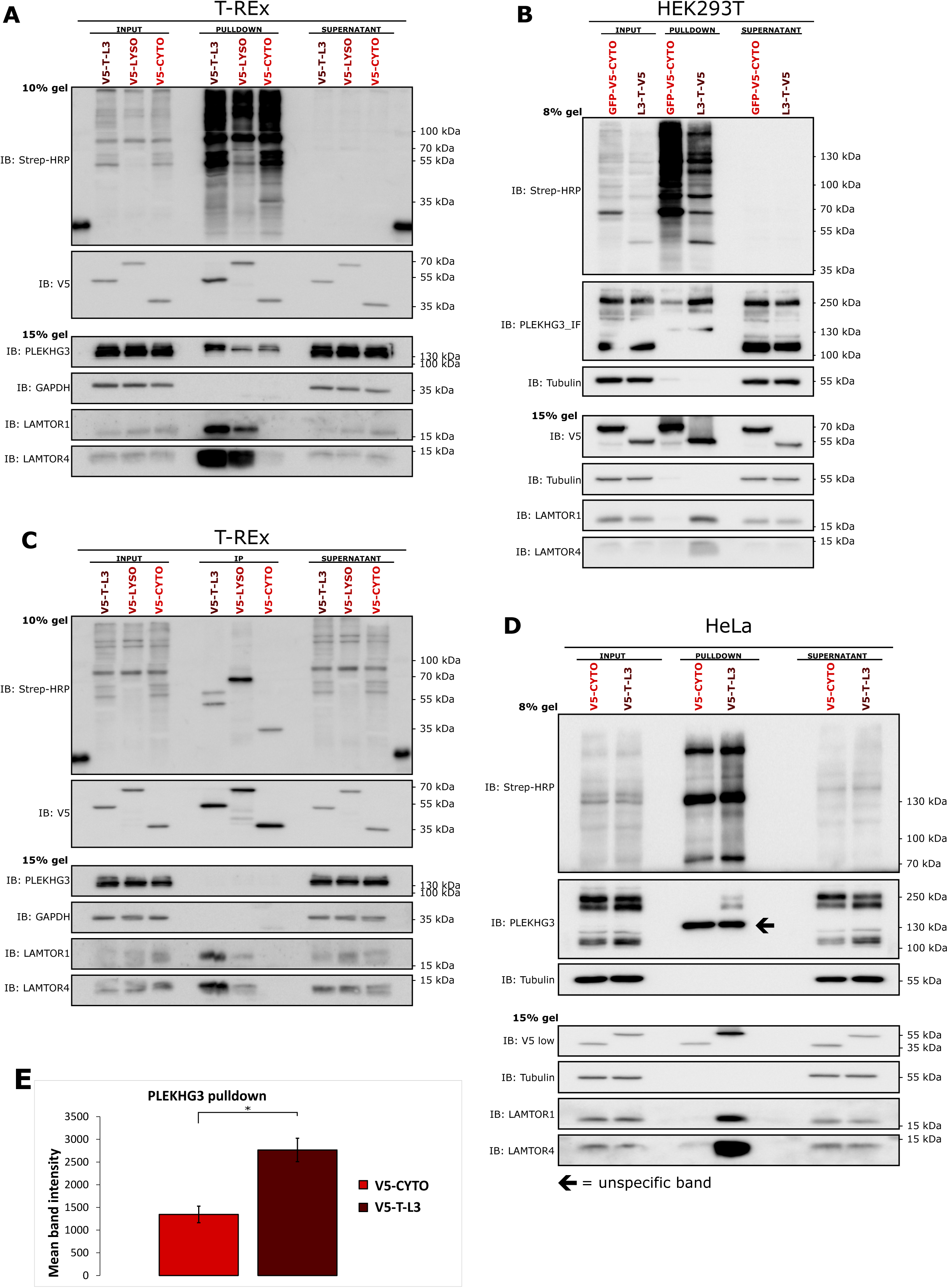
PLEKHG3 is a proximal protein or transient interactor of L3. A-B) Biotinylated proteins were isolated either from HEK293T cells stably expressing L3-T-V5 or GFP-V5-CYTO (left) or from T-REx cells stably expressing V5-T-L3, the V5-LYSO localization control, or the V5-CYTO control (right) and subjected to SDS-PAGE. LAMTOR1 and LAMTOR4 serve as positive controls for direct interaction partners. **C)** V5 immunoprecipitation from T-REx cells expressing V5-T-L3, the V5-LYSO localization control, or the V5-CYTO control were subjected to SDS-PAGE and immunoblotted with a PLEKHG3 antibody to detect interaction. **D)** Representative immunoblot of proteins biotinylated by V5-T-L3 and V5-CYTO. GAPDH serves as a loading control (input and supernatant) and as a specificity control for the pulldowns. The band marked by an arrow is only present in the streptavidin pulldowns but not in the input or supernatant and is therefore considered unspecific. **E)** Quantification of PLEKHG3 bands from immunoblots similar to the one shown in C. Error bars = SEM, n=3. * = p values according to student’s t-test.

**Figure 3:**
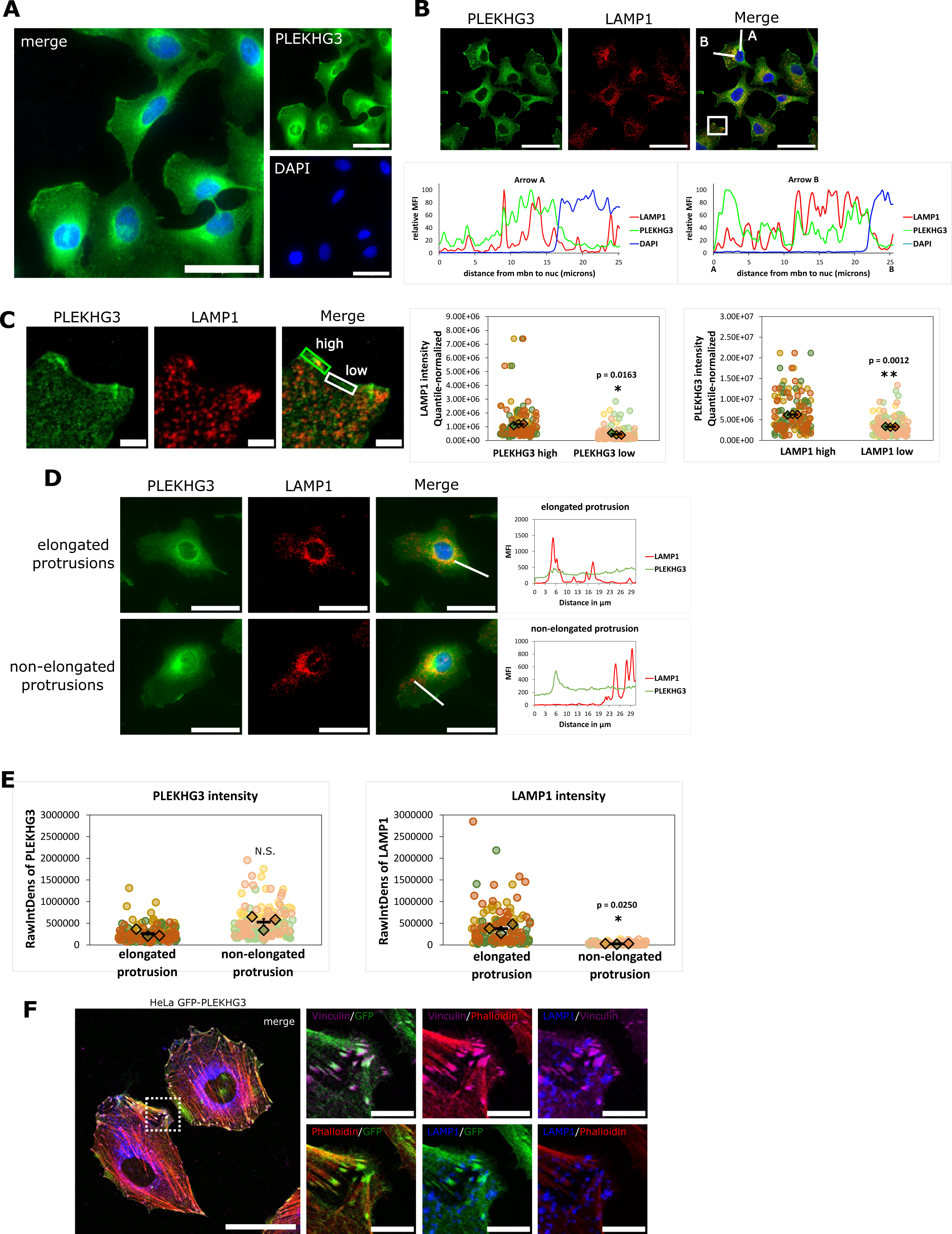
PLEKHG3 and lysosomes colocalize at protrusions in the cell periphery. **A)** Immunofluorescence images of HeLa cells stained for endogenous PLEKHG3 (green) and DAPI (blue). Scale bar = 50 µm. **B)** Immunofluorescence images of HeLa cells stained with PLEKHG3 (red) and LAMP1 (grayscale) antibodies show an enrichment of lysosomes at PLEKHG3-high regions of the PM. White lines are drawn from the extracellular space to the nucleus, either over a PLEKHG3-high region (line A) or over a PLEKHG3-low region (line B) of the PM. The intensity of the PLEKHG3/LAMP1 staining is shown in profile plots below the images; scale bar = 50 µm **C)** Higher magnification of the PM region boxed in white in B. This a representative image showing PLEKHG3-high regions outlined in green, PLEKHG3 low regions in white; scale bar = 5 µm. Images were acquired on Olympus slide scanner. **D)** PLEKHG3 and LAMP1 content in elongated (top panels) or non-elongated protrusions (bottom panels). Endogenous PLEKHG3 is shown in red, LAMP1 in grayscale. Scale bar = 30 µm. The profile plots (right) show PLEKHG3 and LAMP1 intensity along the white lines drawn on the merged images. **E)** Quantification of PLEKHG3 and LAMP1 intensity in elongated vs. non-elongated protrusions. Images in A-D were acquired on Olympus slide scanner. **F)** Confocal immunofluorescence image of HeLa cell stably expressing GFP-PLEKHG3 and immunostained with LAMP1 (lysosomes), Vinculin (FAs) and stained with Phalloidin (F-actin). Scale bar = 50 µm. Insets show indicated channels to better visualize colocalization events. Scale bar = 10 µm. **A-D** show epifluorescence images. In the superplots in **C and E**, dots represent individual data points of each of the three-color coded replicates; diamonds represent the mean of each replicate; black bars represent the mean ± the SEM of three biological replicates; * = p values according to student’s t-test.

**Figure 4:**
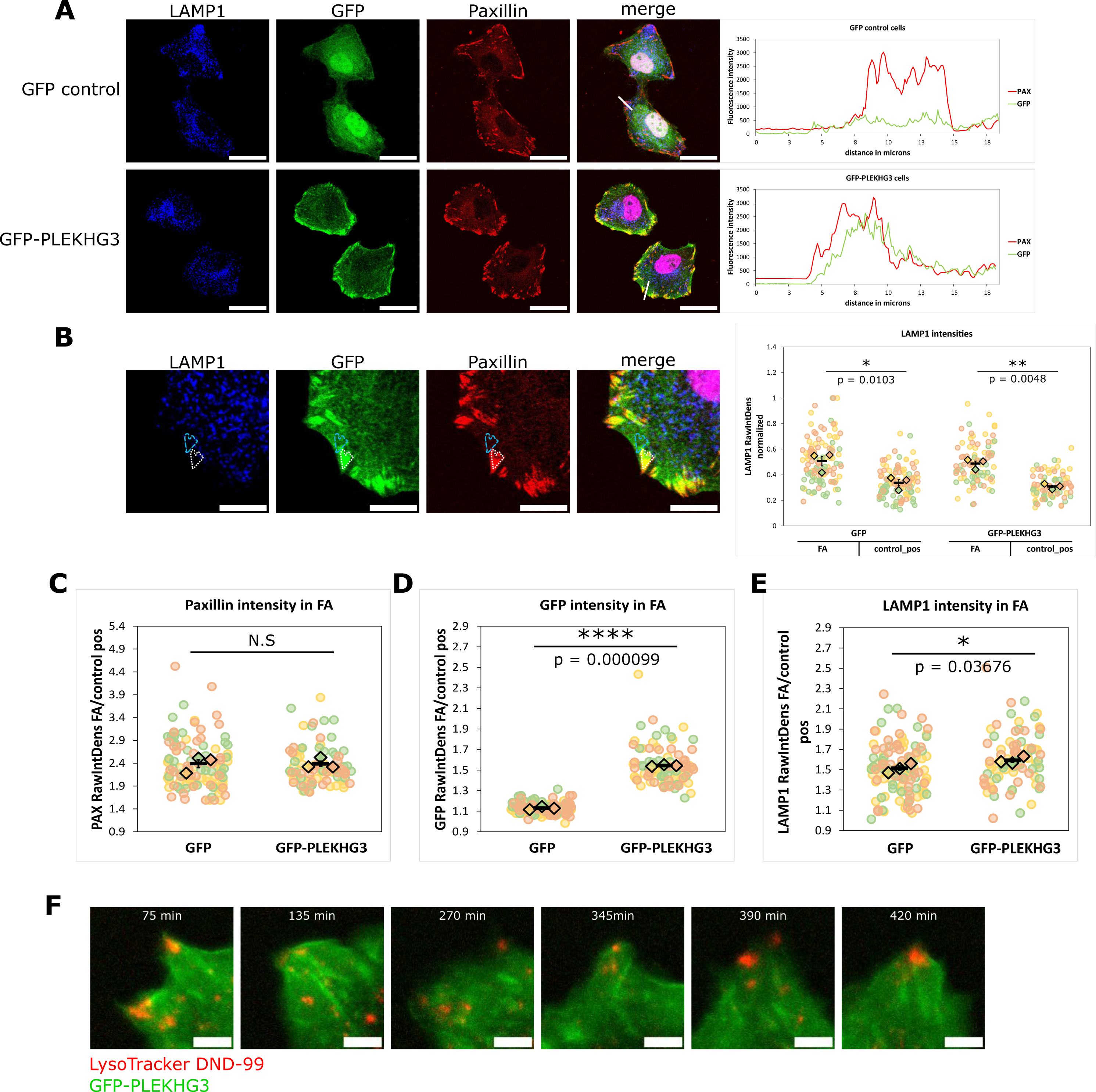
PLEKHG3 accumulates and colocalizes with lysosomes at FA sites. **A)** Confocal images of HeLa cells stably expressing GFP or GFP-PLEKHG3 (green). Cells were stained with Paxillin antibodies (red) to label FAs and with LAMP1 antibodies (grayscale) to label late endosomes/lysosomes. White lines correspond to profile plots on the right. Scale bar = 30 µm. **B)** Confocal images for schematic representation of colocalization analysis. HeLa cells stably expressing GFP-PLEKHG3 were stained with Paxillin and LAMP1 antibodies as described above. PLEKHG3 and LAMP1 intensity was measured in FAs, identified using the Pax channel (dotted white line), and in adjacent, Pax-negative positions of similar shape and area (control positions, broken blue line; see materials and methods section for details) Scale bar = 10 µm. The right panel shows quantification of LAMP1 intensity in paxillin-positive FA versus matched adjacent control regions. **C-E)** GFP/PAX/LAMP1 colocalization. Average intensity of the respective fluorophores in an average of 50 FA per cell. In the superplots, dots represent individual data points of each of three color coded biological replicates of experiments similar to the one shown as immunofluorescent image; diamonds represent the mean of each replicate; black bars represent the mean ± the SEM of the three biological replicates; * = p values according to student’s t-test. **F)** Epifluorescence stills from Movie S5. LysoTracker DND-99 in red and GFP-PLEKHG3 in green show lysosomal movement into cell protrusions. Images were acquired every 15 minutes on a Zeiss Celldiscoverer 7.

**Figure 5:**
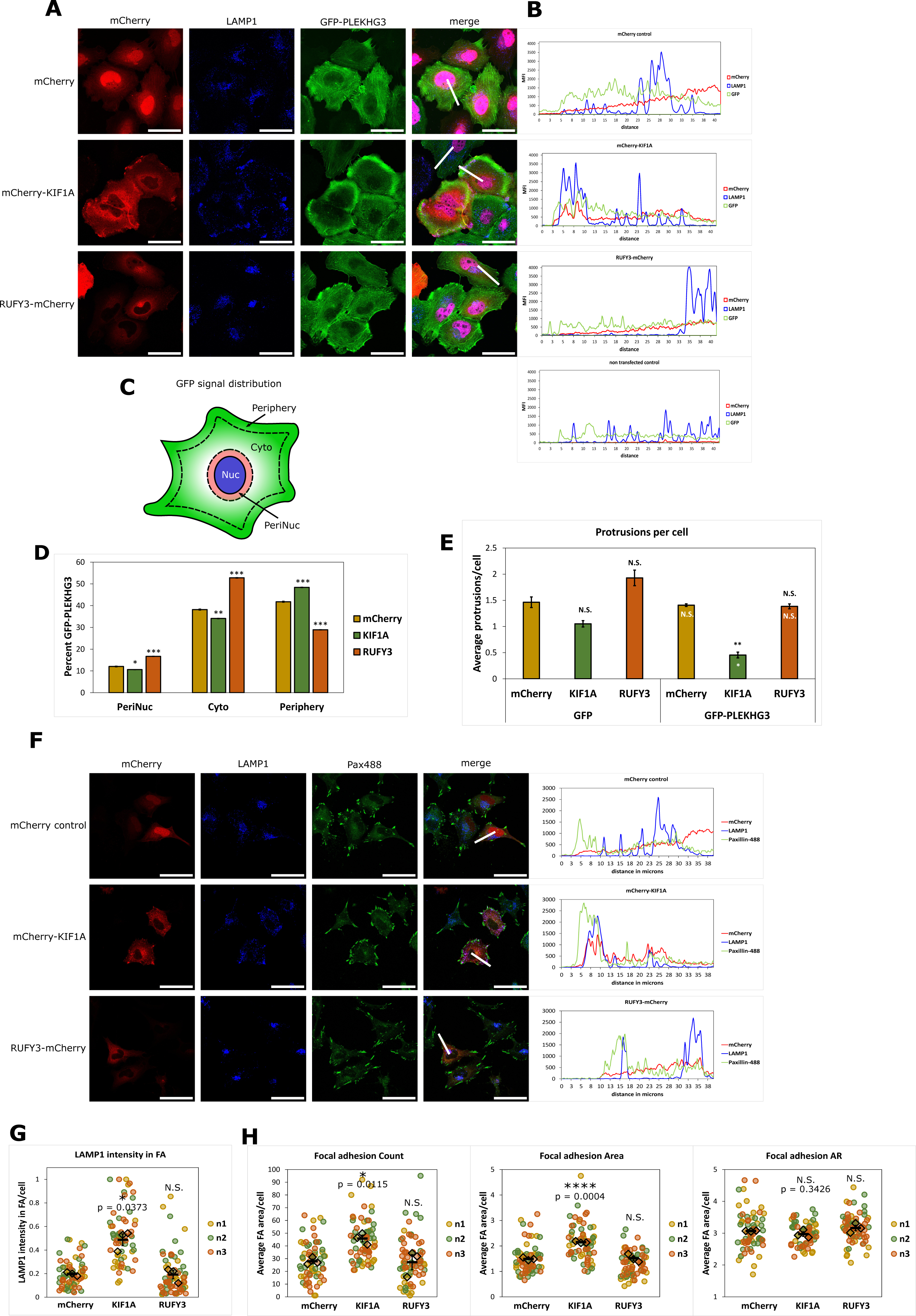
Effect of lysosomal positioning on PLEKHG3 and FAs. A-B) Confocal immunofluorescence images of HeLa cells stably expressing GFP-PLEKHG3, transfected with mCherry constructs (red) and stained with a LAMP1 antibody (grayscale) as well as with DAPI (blue) after fixation. Scale bar = 50 µm. The line plots on the right (B) indicate the distribution of GFP-PLEKHG3, LAMP1 and mCherry along the white lines drawn from the PM to nucleus. **C)** Schematic representation of the distribution analysis. Cells were segmented in Periphery, cytosolic (Cyto) and perinuclear (PeriNuc) regions. GFP intensity was measured in all three segments and the percentage of GFP in each segment was calculated using the intensity of all three segments as 100%. For details see materials and methods. **D)** Quantification of GFP-PLEKHG3 distribution in three independent experiments similar to the one shown in A based on scheme in C. GFP-PLEKHG3 intensity became significantly higher towards the PM. Error bars = SEM. * = p values according to student’s t-test. **E)** Average protrusion number per cell in three independent experiments similar to the one shown in A. Black asterisks = p values according to student’s t-test, comparing the effect of KIF1A or RUFY3 expression with the effect of mCherry. White asterisks in the bars indicate the p values according to student’s t-test comparing GFP-PLEKHG3 and GFP expressing cells. PLE3 = PLEKHG3. **F)** Confocal immunofluorescence images of HeLa cells expressing the indicated mCherry constructs and immunostained for LAMP1 (grayscale) and Paxillin (green). Scale bar = 50 µm. The line plots on the right indicate the distribution of mCherry, LAMP1 and Pax along the white lines drawn from the PM to nucleus. **G)** Lysosomal content in Pax-positive FAs. KIF1A overexpression significantly increases LAMP1 accumulation in FA structures. Superplots show average LAMP1 intensity in FAs per cell. **H)** Morphometric analysis of focal adhesions. KIF1A overexpression increased FA numbers and area. **G-H**, dots represent individual data points of each of three color coded biological replicates of experiments similar to the one shown in F; diamonds represent the mean of each replicate; black bars represent the mean ± the SEM of the three biological replicates; * = p values according to student’s t-test.

### Live cell imaging

Prior to imaging cells were incubated with 60 nM LysoTracker Red DND-99 (L7528, Invitrogen), 50 nM LysoTracker Deep Red (L12492, Invitrogen) or 50 nM LysoTracker Green DND-26 (8783P, Cell Signaling Technology) in imaging DMEM (31053-028; Gibco) without phenol red supplemented with HEPES buffer 25 mM (H0887-100ml; Sigma), 10% FCS and 2% L-glutamine for 30 min or 2 hours at 37 °C in precoated 4- or 8-well ibidi glass bottom chambers. Cells were subsequently imaged under a Live Spinning Disk microscope (Visitron/ Yokogawa CSU-W1-T2) using a Plan Apo Lambda 60x/1.42 NA oil objective at 37 °C, 5% CO_2_. Images were acquired depending on the experiments with 405, 488, 561 and 640 lasers for BFP, GFP/LysoTracker Green, mCherry/LysoTracker Red and LysoTracker Deep Red, respectively on an EM-CCD camera. 5-slice z-stack images were taken every 3 or 4 minutes on multiple positions for a time course of 2-3 hours depending on the experiment. Software used was VisiView. Where indicated, cells were imaged on a Celldiscoverer 7 (Zeiss) using a Plan-Apochromat 50x/1.2 NA water-immersion objective at 37 °C, 5% CO_2_ taking images every 15 min. Images were acquired with a Hamamatsu ORCA Flash4.0 V2 sCMOS camera using the software Zeiss ZEN 3.1. All movies were processed in ImageJ. For presentation purposes z-projections of videos with constant contrast and brightness adjustments in every experiment are shown.

### Immunoblotting

Immunoblotting was performed as follows: Cells were washed three times in ice cold PBS (D8537; Sigma-Aldrich) and lysed in RIPA buffer (50 mM Tris-HCl, pH 7.4, 150 mM NaCl, 1% Triton X-100, 0.5% sodium deoxycholate, and 0.1% SDS) supplemented with cOmplete™, Mini, EDTA-free protease-inhibitor cocktail (40694200; Roche) on ice. Protein concentrations of lysates were measured using a Pierce™ BCA protein assay kit (23225; Pierce), and equal amounts of protein (30 µg) were subjected to SDS-PAGE and immunoblotted. The following primary antibodies were incubated on membranes overnight at 4°C: rabbit anti-LAMTOR1 (1:1000; #8975), rabbit anti-L3 (1:1000; #8168), rabbit anti-LAMTOR4 (1:1000; #13140), rabbit anti-LAMTOR5 (1:1000; #14633), rabbit anti-RAPTOR (1:1000; #2280), rabbit anti-AMPK (1:1000; #2532), rabbit anti-RagC (1:1000; #9480), rabbit anti-GFP (1:1000; #2956), rabbit anti-V5 (1:4000; #13202) all from Cell Signaling Technology; rabbit anti-PLEKHG3 (1:1000; PA5-66764), Streptavidin-HRP (1:2500; 434323) were from Invitrogen; rabbit anti-PLEKHG3 (1:1000; HPA063455), mouse anti-α-Tubulin (1:10,000; T5168) were from Sigma-Aldrich and other antibodies were rabbit anti-mCherry (1:2000; 26765-1-AP; Proteintech),rabbit anti-RUFY3 (1:1000; LS-C681236; LSBio), rabbit anti-PLEKHG3 (1:1000; AP11167b; Abcepta), rabbit anti-GAPDH (1:10,000; ABS16) from Merck and rabbit anti-TMEM192 (1:100; ab236858; Abcam). Secondary antibodies (Jackson ImmunoResearch) were all conjugated to horseradish peroxidase and incubated at a dilution of 1:5000 for 1h at room temperature. SuperSignal™ West Pico PLUS ECL reagent (34580; Thermo-Fisher Scientific) was used to visualize chemiluminescence on a ChemiDoc imaging system (Bio-Rad Laboratories). Quantification of Immunoblots was performed where indicated using ImageJ (version 1.41; National Institutes of Health).

### Pulldowns and immunoprecipitations

Streptavidin-pulldowns were performed as follows: All cells were incubated with 500 µM Biotin (Sigma-Aldrich; B4501) for 10 min in medium containing 10% FCS. Subsequently, cells were washed thrice with ice cold 1x phosphate buffered saline (PBS) (TMS-012; Sigma-Aldrich). HeLa cells were rinsed in the dish while kept on ice and HEK293T or T-REx cells were carefully scraped from the plate and spin-washed in 15 ml falcon tubes in a table-top centrifuge at 300 rcf, 4 °C for 5 min. After the last washing step supernatant was aspirated and cells were lysed in RIPA lysis Buffer (50 mM Tris-HCl, pH 7.4, 150 mM NaCl, 1% Triton X-100, 0.5% sodium deoxycholate, and 0.1% SDS) supplemented with cOmplete™, Mini, EDTA-free protease-inhibitor cocktail (40694200; Roche). After protein concentration determination by BCA protein assay kit (23227; Thermo-Fisher Scientific) 300 µg protein in 600 µl total volume was incubated with 30 µl prewashed Streptavidin magnetic beads (88817; Pierce) at 4 °C overnight on a rotating wheel. For input samples 5-15 µg were used. After incubation 5-15 µg of supernatant were drawn from samples. Beads were washed three times with RIPA lysis buffer and eventually proteins were eluted from beads in 2x sample buffer by vigorously vortexing at room temperature. Samples were then boiled for 7 min at 95 °C and subsequently subjected to Immunoblot analysis.

Immunoprecipitations were performed in a similar manner: In brief, cells were also washed three times in ice cold PBS (rinse for HeLa cells, spin-wash for T-REx cells) and lysed in ice cold IP lysis buffer (20 mM TRIS-HCl, 137 mM NaCl, 1 mM CaCl_2_, 1 mM MgCl_2_, 1% Triton X-100, 2 mM sodium-fluoride, 2 mM Phenylmethylsulfonylfluoride, 1 mM sodium-orthovanadate) supplemented with cOmplete™, Mini, EDTA-free protease-inhibitor cocktail (40694200; Roche). After protein concentration measurement the 300 µg of protein were incubated with 30 µl of anti-V5 tag magnetic beads (MBL International Corporation; M167-11) overnight at 4 °C on a rotating wheel. Further sample processing was equal to the above explained.

### Treatments for TurboID screen

Amino acid starvation (EBSS) was performed by incubation in EBSS medium (14155048; Gibco) supplemented with 200 mg/L CaCl_2_ + 200 mg/L MgCl_2_ for 30 min. Restimulation (HF) was achieved by exchanging starvation medium with full medium containing 10% FCS for 10 min. FM condition was performed by incubation with medium containing 10% FCS for 50 min prior to lysis.

### TurboID-based proximity-dependent labeling

pCDNA5_GOI FRT/TO (FLP recombination target/TetO) plasmids were constructed using the NEBuilder assembly tool (version 2.7.1; New England Biolabs) and generated by Gibson assembly according to the manufacturer’s instructions (New England Biolabs; E2611S). In brief, pCDNA5 FRT/TO vector was cut open by restriction digestion with XhoI and BamHI. V5-TurboID-L3 insert was generated by PCR amplification from pCW57.1-V5-TurboID-L3 (inhouse) plasmid using the primers 5’-taagcttggtaccgagctcggatccAGCCACCATGGGCAAGCC-3’ (forward) and 5’-tttaaacgggccctctagactcgagTTAAGACACTTCCACAACTTTTATCAGTTCTTCAAATAAC -3’ (reverse). V5-TurboID-TMEM192 insert was generated by separately PCR amplifying V5-TurboID insert from pCW57.1-V5-TurboID-L3 plasmid (inhouse) using the primers 5’-taagcttggtaccgagctcgGCCACCATGGGCAAGCCC-3’ (forward) and 5’-cccccgccgcCTGCAGCTTTTCGGCAGACC-3’ (reverse) and the TMEM192 insert from pLJC5-TMEM192-2xFLAG (102929; Addgene) using the primers 5’-aaagctgcagGCGGCGGGGGGCAGGATG -3’ (forward) and 5’-tttaaacgggccctctagacTTACGTTCTACTTGGCTGACAGCCCAGGTC-3’ (reverse). The V5-TurboID-NES insert was generated by PCR amplification from V5-TurboID-NES_pCDNA3 (107169; Addgene) using the primers 5’-taagcttggtaccgagctcggatccAGCCACCATGGGCAAGCC-3’ (forward) and 5’-tttaaacgggccctctagactcgagTTAGTCCAGGGTCAGGCGC-3’ (reverse). Inserts were mixed with the linearized pCDNA5 FRT/TO vector and fused by Gibson assembly. Stable T-REx cells were generated by using the Flp-In™ T-REx™ Core Kit (K650001; Invitrogen) according to the manufacturer’s instructions. FlpIn T-REx host cells were transfected with pcDNA5-V5-TurboID-L3 FRT/TO, pcDNA5-V5-TurboID-TMEM192 FRT/TO or pcDNA5-V5-TurboID-NES FRT/TO and pOG44 at a ratio of 400 ng pcDNA5 FRT/TO to 2 µg pOG44. Afterwards, cells were selected in 100 µg/ml hygromycin B (H3274; Sigma-Aldrich). Cells were maintained in DMEM with 10% doxycycline-free FBS, 100 µg/ml hygromycin, and 15 µg/ml blasticidin at 37°C, 5% CO_2_. Expression of constructs close to endogenous levels was induced by incubation with 1 ng/ ml doxycycline for pcDNA5-V5-TurboID-L3 FRT/TO and pcDNA5-V5-TurboID-NES FRT/TO or 0.1 ng/ ml doxycycline for pcDNA5-V5-TurboID-TMEM192 FRT/TO for 24 hours. Proximity labelling was carried out as follows: Cells were incubated with 500 µM biotin (in DMSO) full medium for 15 min at 37°C, 5% CO_2_. The reaction was stopped by transferring plates on ice and medium was decanted. Cells were immediately spin-washed three times with cold PBS and subsequently lysed in RIPA buffer supplemented with cOmplete™, Mini, EDTA-free protease-inhibitor cocktail (40694200; Roche) on ice. Cell lysates were transferred to low-binding microcentrifuge tubes (022431081; Eppendorf) and 1 mg protein in 1 ml total volume was incubated with 200 µl prewashed Streptavidin magnetic beads (88817; Pierce) at 4 °C overnight on a rotating wheel. The beads were washed three times with RIPA buffer and five times with detergent-free washing buffer (50 mM Tris-HCl, 50 mM NaCl, pH 7.4). Beads were kept on ice until further processing.

### Sample preparation for mass spectrometry analysis

Beads were resuspended in 50 µL 1 M urea, 50 mM ammonium bicarbonate. Disulfide bonds were reduced with 2 µL of 250 mM dithiothreitol (DTT) for 30 min at room temperature before adding 2 µL of 500 mM iodoacetamide and incubating for 30 min at room temperature in the dark. Remaining iodoacetamide was quenched with 1 µL of 250 mM DTT for 10 min. Proteins were digested adding 300 ng trypsin (Trypsin Gold, Promega) followed by incubation at 37°C overnight. The digest was stopped by addition of trifluoroacetic acid (TFA) to a final concentration of 0.5 % and the supernatant was transferred to a fresh tube. The peptides were cleaned up according to the SP2 protocol by Waas et al. [3].

### Liquid chromatography - mass spectrometry analysis

Peptides were separated on an Ultimate 3000 RSLC nano-flow chromatography system (Thermo-Fisher Scientific), using a pre-column for sample loading (Acclaim PepMap C18, 2 cm × 0.1 mm, 5 μm, Thermo-Fisher Scientific), and a C18 analytical column (Acclaim PepMap C18, 50 cm × 0.75 mm, 2 μm, Thermo-Fisher Scientific), applying a segmented linear gradient from 2% to 35% over 120 min and finally 80% solvent B (80 % acetonitrile, 0.08 % formic acid; solvent A 0.1 % formic acid) at a flow rate of 230 nL/min. The nano-LC was interfaced online via a nano-spray ion-source to a Q Exactive HF-X Orbitrap mass spectrometer (Thermo-Fisher Scientific). The mass spectrometer was operated in a data-dependent acquisition mode (DDA). Survey scans were obtained in a mass range of 375-1500 m/z, at a resolution of 120k at 200 m/z and an AGC target value of 3E6. The 8 most intense ions were selected with an isolation width of 1.6 m/z, at a target value of 1E5 with a max. IT of 250 ms, and fragmented in the HCD cell with 28% normalized collision energy. The spectra were recorded at a resolution of 30k. Peptides with a charge of +2 to +6 were included for fragmentation, the peptide match and the exclude isotopes features were enabled. Selected precursors were dynamically excluded from repeated sampling for 30 seconds.

### Database search

MS raw data were analyzed using the MaxQuant software package (version 1.6.0.16, Tyanova et al., 2016) searching the Uniprot *Homo sapiens* reference proteome (www.uniprot.org), the sequences of the protein constructs as well as a database of common contaminants. The search was performed with full trypsin specificity and a maximum of two missed cleavages at a protein and peptide spectrum match false discovery rate of 1%. Carbamidomethylation of cysteine residues was set as fixed, oxidation of methionine and acetylation of the protein N-terminus as variable modifications. For label-free quantification the “match between runs” feature and the LFQ function were activated - all other parameters were left at default.

### Proteomics data analysis

The output from MaxQuant was analyzed using amica [4]. Proteins from reverse database identifications, potential contaminant proteins and proteins only identified by a modified peptide were removed. Proteins with at least 2 Razor + unique peptides, at least 3 MS/MS counts, and valid values in 3 replicates in at least one group were considered quantified. LFQ intensities of quantified proteins were log2-transformed and quantile normalized. Intensities with missing values were imputed from a normal distribution downshifted 1.8 standard deviations from the mean with a width of 0.3 standard deviations. Differential expression analysis was performed with DeqMS (version 1.10.0) [5].

### Over-Representation Analysis

Enriched proteins (log2 FC > 0 and padj ≤ 0.05) being both significantly enriched in the pairwise group comparisons L3/Cyto full medium and Lyso/Cyto full medium were used as an input for an over-representation analysis using gprofiler2 (version 0.2.1) [6] with the database Ensembl 104, Ensembl Genomes 51 (database built on 2021-05-07). The three branches of the Gene Ontology (GO:CC, GO:MF, GO:BP) [7,8] were used as source databases for gprofiler2, excluding electronic GO annotations.

### Network analysis

For the N-terminally tagged L3 bait (V5-T-L3), enriched proteins (log2 FC ≥ 1.49 and padj ≤ 0.05) were considered from all three conditions (pairwise group comparisons L3/Cyto in full medium, EBSS and EBSS+full medium). To remove unrelated nuclear proteins from this list of preys, proteins annotated with the term ‘nucleus’ (GO:0005634) that were not significantly enriched in the lyso control in at least one condition (log2 FC ≥ 1.49 and padj ≤ 0.05) were removed. The overlap of these preys in the three conditions was plotted (Figure S1J + K) using the eulerr package (version 6.1.1), and the pheatmap package (version 1.0.12) using colors from ColorBrewer [9]. For the C-terminally tagged L3 bait (L3-T-V5), enriched proteins (log2 FC ≥ 1.49 and padj ≤ 0.05) were filtered in the same way to remove nuclear proteins.

Cytoscape (version 3.8.2) [10] was used to create functional networks (Figure 1D + F, Figure S1D + I) of significant preys. Gene names from enriched proteins were loaded into the stringApp (version 1.7.1) [11] which returned a full STRING network (v11.5) [12] with a default confidence score cutoff of 0.4. Networks were visualized with the yFiles (v1.1.2) organic layout algorithm.

### Quantification and statistical analysis

All image analysis was performed in Fiji distribution of ImageJ [13]. Where indicated by white lines, the corresponding plots show the intensity profile of the respective channels along the length of the line. Profile plots were generated using the Plot Profile function in Fiji. Quantification of PLEKHG3 knockdown in immunofluorescence (IF) images: Single cells were segmented based on their combined PLEKHG3 and GFP channels by thresholding. From the resulting cell masks the integrated density (RawIntDens) was measured for both, GFP and PLEKHG3 channels after background subtraction. Superplots represent average values of 11 cells per condition from three technical replicates. Analysis of PLEKHG3 KO: A minimum of 50 cells per replicate per condition was segmented out from slide scanner images and analyzed for integrated PLEKHG3 signal density and compared using a 2-way ANOVA. Colocalization of Phalloidin with GFP and GFP-PLEKHG3 (Figure S3C) or LAMP1 with LAMTOR4 (Figure S3F) was performed on background subtracted images using the JACoP plugin [14]. Overlap of endogenous PLEKHG3 with LAMP1 and vice versa was analyzed from background subtracted slide scanner images. High intensity and adjacent low intensity regions were selected as ROIs in both channels and measured by RawIntDens. Similarly, the content of PLEKHG3 and lysosomes in elongated vs non-elongated protrusions was quantified by selecting them as ROIs and measuring RawIntDens from both channels. Colocalization analysis of FA structures with GFP, GFP-PLEKHG3, LAMTOR4 and LAMP1 was performed as follows: FA structures from single cells were defined as ROIs based on the Paxillin staining by segmentation which involved thresholding and particle size filtering. Control regions were defined by translocating the FA ROIs to adjacent regions. RawIntDens was measured for all channels of the respective experiment in all channels. Morphometric analysis was achieved by same FA segmentation as described above. Shape descriptor tools from Fiji (Area, Circularity, Aspect ratio) were applied on all FA ROIs. Morphometric analysis of cell shape was performed by segmenting single cells based on their GFP staining. Resulting cell masks were analysed for the above-mentioned shape descriptors. In live cell imaging analysis, the shape descriptors were measured for all time points and then averaged over time. GFP-PLEKHG3 and BFP-LifeAct colocalization (Figure S6C) was measured over time by Fijis Coloc2 Plugin. We defined protrusive activity as the combination of forming and retracting protrusions of a cell over time. Protrusions were counted for each cell individually and indicated by yellow and cyan arrows representing forming and retracting protrusions, respectively. Quantification was performed on ≥ 14 cells per condition and quantified by non-parametric Mann-Whitney U or Kruskal-Wallis test, based on number of compared groups with a post-hoc p-value adjustment according to Bonferroni to account for unequal variance and non-gaussian distribution. GFP and GFP-PLEKHG3 distribution in the cell was quantified by dividing the cells into peripheral, cytoplasmic and perinuclear regions. Perinuclear region was defined by enlarging the nucleus’ outline based on DAPI staining and by 4 µm and subtraction of nucleus outline. Similarly, the cell outline was defined based the GFP channel which was subtracted by the outline shrunk by 4 µm. The cytosolic region was therefore defined as the whole cell of which peripheral and perinuclear regions were subtracted. For all regions RawIntDens was measured. Lysosomal distribution was performed based on an adapted Fiji macro from Filipek et al. [15]. In brief, the macro defines the center of mass of the nucleus based on DAPI staining and from there creates concentric annular ROIs with increasing radii. RawIntDens is measured in background subtracted LAMP1 channel. Quantification was performed on ≥ 14 cells (confocal images) or ≥ 50 cells (slide scanner images) per condition from three biological replicates. Quantification of immunoblots was performed in Fiji. Two-tailed, paired Student’s t tests were performed on experimental data from at least three independent experiments. Asterisks indicate the following significances: * = p < 0.05; ** = p < 0.01; *** = p < 0.001; **** = p < 0.0001.

## Introduction

Lysosomes are acidic organelles recognized for their role in degradation of proteins, nucleic acids and other macromolecules to maintain cellular metabolic homeostasis. Beyond that, lysosomes are highly dynamic structures that constantly modulate their size and cellular distribution and facilitate contact and fusion events with other organelles such as the endoplasmatic reticulum (ER). The lysosome limiting membrane serves as a signaling platform containing a wide repertoire of residents, such as transmembrane and membrane-associated proteins, protein complexes that dynamically recruit cytosolic proteins involved in signaling and metabolic processes as well as communication to other organelles [16,17].

One of these resident protein complexes is the late endosomal/lysosomal adaptor and MAPK and mTOR activator (LAMTOR) complex, also known as Ragulator. This pentameric complex is essential for recruiting the small Rag GTPases essential for mTORC1 regulation. Structurally, the LAMTOR complex consists of two heterodimers LAMTOR2 (p14) /LAMTOR3 (MP1) and LAMTOR4 (C11orf59)/LAMTOR5 (HBXIP) held together by the belt-like, myristoylated LAMTOR1 (p18) subunit that anchors them to the lysosomal surface [18]. The LAMTOR complex integrates mTORC1 activation [19], MEK/ERK signaling [20], AMPK signaling [21], lysosomal size regulation [22,23] actomyosin contractility in immune cells [24] and lysosome positioning [15,25,26].

Spatial positioning of the lysosome dictates physico-chemical properties, including luminal pH, content and activity of digestive enzymes [27,28]. Peripheral accumulation of lysosomes has been correlated with increased mTORC1 activation [29–31] and more specifically with the association of lysosomes with focal adhesion (FA) sites [32]. Peripheral lysosomes have also been implicated in fusion processes occurring at the plasma membrane (PM), such as antigen presentation, membrane repair and cellular clearance by lysosomal exocytosis, through which they release intracellular content into the extracellular space [17,33–36]. Interestingly, peripheral localization of lysosomes and their subsequent exocytosis was shown to be crucial for cancer cell migration [37]. To reach the PM, lysosomes have to transition from the microtubule tracks, on which they move through the cell, to the cortical actin mesh [38,39]. It has been shown that lysosomes use myosin motor proteins to move along actin filaments in a manner similar to their kinesin- or dynein-directed movement on microtubule tracks [40,41]. The interaction with the dense cortical actin network causes transient retention of lysosomes in the periphery of the cell [40,42]. Recently, a study revealed a significant role of peripheral lysosome accumulation for collective cell migration and leader cell emergence [43].

While a vast array of approaches has sought to define the lysosomal proteome, [44–48], the identification of transient interactors or vicinal proteins has remained a challenge. Recently, utilization of proximity-dependent labeling screens [49–52] has led to the identification of previously unknown lysosomal and lysosome-associated proteins. However, these approaches were either limited by the ligase used due to long labeling times (BirA*), the requirement of cytotoxic reagents (APEX) or by the baits used. In this study, we utilized a TurboID proximity-dependent labelling screen using LAMTOR3 as bait to identify cytoplasmic proteins recruited to the lysosomal surface. One of the main hits in the LAMTOR3 (L3) proxisome was PLEKHG3, a protein that contributes to cortical actin remodeling by binding selectively to newly polymerized actin at the cell’s leading edge. PLEKHG3 acts as a GEF for Rac1 and CDC42 downstream of PI3K, inducing positive feedback on actin polymerization and promoting Rac-mediated membrane ruffling downstream of RHOG. [53,54]. We demonstrate that PLEKHG3 colocalizes with lysosomes at focal adhesion and protrusion sites, and that the localization and function of this protein – and consequently, overall cell motility – are fundamentally dependent on lysosomal dynamics.

## Results

### Proximity-dependent labelling reveals an interaction between LAMTOR3 and the actin remodeling network

The pentameric LAMTOR/Ragulator complex has been established as a signaling hub on the lysosomal membrane that orchestrates metabolic and growth factor signaling. To interrogate proteins recruited to the limiting membrane of lysosomes, we employed unbiased mass spectrometry (MS)-coupled proximity-dependent labeling (Figure 1A) using LAMTOR3 (L3) as bait. Therefore, we generated stable (dox-inducible) HEK293T cell lines, expressing LAMTOR3 fused to a C-terminal TurboID-V5 tag (L3-T-V5) and GFP-NES (nuclear export signal; GFP-T-V5) as control lines. Bait expression levels were matched (Figure S1A-B). Immunofluorescence confirmed the predominantly lysosomal localization of the L3-T-V5 bait and its colocalization with the lysosomal-associated membrane protein 1 (LAMP1) (Figure S1C). Analysis of prey proteins specifically enriched by the L3 bait yielded an interaction network of known L3 binding partners including the other LAMTOR complex subunits, parts of the BORC complex, the neutral amino acid transporter SLC38A9 as well as several proteins involved in regulation of mTORC1 activity. While we were able to validate well-established interactors, the dataset importantly recovered previously uncharacterized proteins enriched by L3 (Figure S1D).

To ensure comprehensive and reliable mapping of LAMTOR3 interactors, we performed an ortholog screen using N-terminal tagged LAMTOR3 (V5-T-L3), complementing the C-terminal tagged screen (V5-TurboID fused to NES serves as a cytosolic specificity control: V5-CYTO; see Figure S1E). Given the predominant lysosomal localization of LAMTOR3 baits (Figure S1C and F), we generated a localization control for lysosomal biotinylation (V5-LYSO; V5-TurboID fused to the lysosomal transmembrane protein TMEM192) (Figure S1E). Using the doxycycline-inducible Flp-In™ T-REx™ system (Invitrogen™) we achieved close to endogenous, homogenous expression of the bait proteins (Figure S1G). To assess the dynamic nature of protein dynamics, we exposed cells to three different states: full medium (FM), starvation (EBSS, growth factor + amino acid starvation) and restimulation (EBSS + 10 minutes FM), to discover interactions induced by these conditions (Figure S1H). Our dual-screening approach demonstrated robustness, corroborating findings of the previous screen. Examples include the LAMTOR subunits (LAMTOR1-5), Rag GTPases and RAPTOR as a representative subunit of the mTORC1 complex, parts of the BORC complex, as well as a subunit of the vATPase (Figure S1I). Indeed, the majority of the hits (∼64% for V5-T-L3 and ∼67% for L3-T-V5) were identified in both screens, irrespective of the differences in the experimental systems used (specific vs random integration, homogenous vs heterogenous expression; cell line specific differences; ligase positioning in the complex; cytosolic control baits), strengthening confidence in the robustness and reliability of the screens (Figure 1B). Interestingly some preys showed bait specificity, most prominently GATOR1 (labeled by L3-T-V5 but not by V5-T-L3 bait). In turn, only V5-T-L3 was able to label the GATOR2 complex (Figure 1B-C) as well as several lysosomal proteins, indirect or transient interactors not identified in previous studies (Figure 1D) (reviewed in [55–57]). Finally, L3-T-V5 labeled subunits of the BORC missed by V5-T-L3, yielding a more complete picture of L3’s interactors (Figure 1D). Somewhat surprisingly, the L3 proxisome was very similar in full-medium, starved, and restimulated cells (Figure S1J-K).

Comparison of the proteins biotinylated by V5-T-L3 and by the V5-LYSO localization control, respectively, revealed a strong overlap, particularly among proteins found in the proximity of the lysosomal membrane. Notably, a distinct subset of hits was specifically biotinylated by V5-T-L3 and could not be detected among the V5-LYSO preys or the V5-CYTO control preys (see Figure S1K).

Overrepresentation analysis (ORA) of the V5-T-L3 interactome identified “molecular adaptor activity” as a primary molecular function, with the lysosomal and endosome membranes as ranking as top-scored cellular component. mTORC1 and autophagy regulation were at the top of the biological processes (Figure 1E). While these categories were expected, we also identified a specific subset of proteins involved in actin remodeling, including the GO-terms “GEF activity and actin filament binding” and “localization to the actin cytoskeleton” (Figure 1E-F). Most of these hits were significantly enriched although not very abundant (Supplementary Table 1), the RhoGEF PLEKHG3 emerged as a top hit in both screens (Figure 1C and F). PLEKHG3 is a protein consisting of a Dbl-homology (DH) domain to execute its RhoGEF activity, a Pleckstrin homology (PH) domain important for its binding to phosphoinositides at the PM, and an actin binding domain (ABD). PLEKHG3 has been described to colocalize with actin filaments, more specifically accumulating at sites of membrane remodeling such as the leading or the trailing edges. Its identification in our screens suggests a novel role for lysosomes in actin remodeling, possibly mediated by L3. [53,54].

### PLEKHG3 is a vicinal protein of LAMTOR3 that colocalizes with peripheral lysosomes at cell protrusions

Based on the identification of PLEKHG3 as a high-confidence proximal interactor, we proceeded to characterize its endogenous localization and relationship to L3. We first validated the specificity of the PLEKHG3 antibody using a dual siRNA and CRISPR/Cas9 approach. Immunoblotting displayed multiple bands with molecular weights (MW) whose intensity dropped significantly upon siRNA-based knockdown of PLEKHG3 both, in HeLa and in HEK293T cells (Figure S2A-B). Thus, the antibody recognized PLEKHG3, although as previously described [54], the molecular weight of the protein in SDS-PAGE is much higher than the 134 kDa calculated from the amino acid sequence of the protein (Figure S2B).

Immunofluorescence analysis of the same PLEKHG3 antibody revealed colocalization of endogenous PLEKHG3 with Phalloidin in HEK293T cells which was selectively reduced at the plasma membrane upon PLEKHG3 KD (Figure S2C). Confirming the high molecular weight of endogenous PLEKHG3 on the immunoblot (Figure S2B), expression of exogenous GFP-PLEKHG3 also revealed a much higher molecular weight of this protein on SDS-PAGE than the predicted 164 kDa (Figure S2D). Furthermore, colocalization was shown with the exogenous GFP-PLEKHG3 signal, and a significant drop in both the PLEKHG3 and the GFP signal upon PLEKHG3 knockdown (Figure S2E-F). Together, the experiments validated the PLEKHG3 reagents.

CRISPR/Cas9-based ablation of endogenous PLEKHG3 also significantly reduced the high MW band indicated by PLEKHG3 staining on immunoblot, confirming the results generated by the PLEKHG3 knock down (Figure S2G). Also, immunofluorescence analysis of PLEKHG3 KO cells revealed a significant drop in overall PLEKHG3 intensity compared to WT cells (Figure S2H-I). Notably, while the PLEKHG3 signal in the periphery of the cells was abolished upon PLEKHG3 KO, a diffuse perinuclear stain persisted after siPLEKHG3 and in PLEKHG3 KO (Fig. S2C, S2H), indicating non-specific antibody signal; analyses therefore focus on the specific cortical/FA-associated PLEKHG3 signal.

To validate the results obtained by MS, we isolated biotinylated proteins from T-REx and HEK293T cells. We found that PLEKHG3 specifically enriched in streptavidin pulldowns from cells expressing L3 baits (L3-T-V5 or V5-T-L3) compared to their respective controls (Figure 2A-B). In contrast, PLEKHG3 was not detected in V5 co-immunoprecipitates from cells expressing V5-T-L3 or in the V5-CYTO and V5-LYSO controls (Figure 2C). These results suggest that PLEKHG3 is a transient interactor or vicinal protein, rather than a stable, directly binding partner of L3.

We next investigated whether the interaction occurs at a specific subcellular site. Employing HeLa cells – that offer superior morphology for imaging – we validated that V5-T-L3 biotinylated PLEKHG3 at nearly double the efficiency in contrast to the control cytosolic decoy (Figure 2D-E). Together with lack of PLEKHG3 labeling by the TMEM192 localization control and preserved PLEKHG3 localization in L3 KO, these data support spatial vicinality over stable complex formation.

### PLEKHG3 and lysosomes colocalize at the plasma membrane independently of LAMTOR3

Following the identification of PLEKHG3 as a proximal interactor of LAMTOR3, we next characterized its spatial relationship with lysosomes. In HeLa cells, endogenous PLEKHG3 localized to the PM where it colocalized with Phalloidin-labeled actin. This distribution aligns with previous work (Figure 3A and S3A; and [53]), confirming the data from HEK293T cells shown in Figure S2C. Stable expression of GFP-PLEKHG3 in HeLa cells showed a similar localization, but more clearly labeled filamentous actin, particularly cortical actin structures (Figure S3B-C). In addition, co-staining of endogenous PLEKHG3 and LAMP1, used as a late endosomes/lysosomes marker, revealed colocalization right below the PM (Figure 3C). PLEKHG3 accumulation sites showed a strong enrichment of LAMP1 positive vesicles compared to PM regions low in PLEKHG3, and vice versa (Figure 3C). More specifically, we found that PLEKHG3 colocalized more strongly with LAMP1-positive vesicles in elongated membrane structures (Figure 3D-E). Focal adhesion sites, which anchor the intracellular cortical actin network to the extracellular matrix and are remodeled with the help of late endosomes/lysosomes during protrusion formation and cell motility, can also be found in such elongated membrane protrusions (reviewed in [58,59]).

Next, immunofluorescence experiments showed the reported colocalization of endogenous PLEKHG3 (Figure S2C in HEK293T cells, Figure S3A in HeLa cells) and GFP-PLEKHG3 with cortical actin structures and the partial localization of LAMP1-positive vesicles to these structures in correspondence with vinculin-positive focal adhesions (Figure 3F). These findings provide further evidence for the association of PLEKHG3 with the actin cytoskeleton and lysosomal trafficking in our system.

In a next step, we investigated the possible influence of LAMTOR on this process using L3 KO cells (Figure S3D). The knockout of L3 resulted in a strong reduction as well as in a complete dispersion of the LAMTOR4 signal from the lysosome (Figure S3E-F), as expected from earlier reports showing that loss of one LAMTOR subunit causes the disruption of the complex, its delocalization from the lysosome, and a destabilization of the other LAMTOR subunits [60,61]. GFP and GFP-PLEKHG3 constructs were introduced in these cells and expressed at similar levels (Figure S3G). Both, GFP-PLEKHG3 and LAMTOR4 were found enriched in lysosomal structures at paxillin-positive FAs (Figure S3H-I). However, the association of GFP-PLEKHG3 with FA was not impaired in L3 KO cells, nor did the expression of GFP-PLEKHG3 rescue or alter the distribution of LAMTOR4-positive vesicles (Figure S3E-J). Morphometric analysis of FA structures revealed a slight decrease in total FA number in L3 KO cells, while the overall area and aspect ratio remained unaffected (Figure S3K). GFP-PLEKHG3 expression did not affect any of these parameters. Live imaging of cells expressing either GFP or GFP-PLEKHG3 clearly showed that L3 KO cells spread less compared to WT cells, while the expression of GFP-PLEKHG3 had no discernible effect on spreading (Movie S1-4 + Figure S3L).

### Focal adhesions as hubs for convergence of PLEKHG3 and lysosomes

Our data suggests that PLEKHG3 is proximal to LAMTOR3, rather than a stable physical interactor, potentially reflecting a novel association with peripheral lysosomes and focal adhesions. Confirming the data in Figure S3H-I, GFP-PLEKHG3 and LAMP1-positive vesicles were found enriched in FA labeled by paxillin (Figure 4A-E). As expected, based on the enrichment of LAMTOR4 in FAs (Figure S3I) the LAMP1 signal was stronger in paxillin-labeled FA compared to control regions (Plot in Figure 4B).

To further investigate the colocalization of PLEKHG3 with late endosomes/lysosomes, we performed live cell imaging of GFP-PLEKHG3 cells incubated with LysoTracker to label acidic endo-lysosomal structures. Following a single cell over time, we could observe that a subset of lysosomes appears to travel to PLEKHG3 accumulation sites and specifically move into developing protrusions (Figure 4F + Movie S5).

These data are consistent with the hypothesis that lysosomes moving in an anterograde manner reach FA sites [59] where they encounter PLEKHG3 and successively travel in PLEKHG3-induced protrusions.

### Lysosomal trafficking impacts PLEKHG3 localization

As depicted in Figure 4F (and Movie S5), lysosomes traveled into PLEKHG3 accumulation sites, raising the possibility that loss of PLEKHG3 would impair lysosomal outward trafficking. However, CRISPR/Cas9-mediated ablation of PLEKHG3 in HeLa cells showed no effect on lysosomal distribution compared to WT cells (Figure S4A). Similarly, the morphometric parameters (cell area, circularity or aspect ratio) of WT and KO cells were comparable (Figure S4B-C).

Thus, PLEKHG3 ablation did not alter lysosomal trafficking. To determine whether lysosomal trafficking affected the subcellular localization of PLEKHG3, we manipulated lysosomal localization using RUFY3-mCherry to promote lysosomal inward transport [28,62], and mCherry-KIF1A to drive lysosomal centrifugal dispersion [63]. Overexpression of mCherry-KIF1A in GFP-PLEKHG3 expressing stable cell lines shifted the lysosomal localization from the perinuclear region to the cell periphery and induced a concentration of GFP-PLEKHG3 in the cell periphery (Figure 5A-B, profile plots in B; Figure S5B-C). To quantify this effect, we segmented the cytoplasm into 3 areas (perinuclear = PeriNuc, cytoplasm = Cyto, Periphery) and calculated the percentage of PLEKHG3 in each segment by setting the intensity measured in all three segments to 100 % (Figure 5C). While in RUFY3-mCherry the maximum GFP-PLEKHG3 intensity was found in the cytoplasm, in mCherry-KIF1A expressing cells there was a clearly detectable increase towards the periphery (Figure 5D). As a control, the expression of mCherry, mCherry-KIF1A, or RUFY3-mCherry had no effect on the localization of GFP (Figure S5D-E). The effect of the peripheral accumulation of lysosomes on PLEKHG3 distribution could be confirmed by visualizing endogenous PLEKHG3 in cells transiently expressing KIF1B (Figure S5F-H). LAMP1 accumulation sites showed a slight increase in endogenous PLEKHG3 compared to control regions, and vice versa (Figure S5I-K). Concentration of GFP-PLEKHG3 at the PM by mCherry-KIF1A overexpression, however, did not increase the number of protrusions (Figure 5E), in contrast to the reported positive effect of this protein on actin remodeling [53]. Quantification of protrusions clearly indicated a drop upon overexpression of mCherry-KIF1A compared to mCherry control or RUFY3-mCherry overexpression. This effect, consistently present as a trend in GFP control cells, became strongly significant in GFP-PLEKHG3 expressing cells. Concentration of lysosomes around the nucleus by overexpression of mCherry-RUFY3 had no significant effect on protrusion number compared to control conditions (Figure 5E).

In terms of morphology, the cells expressing mCherry-KIF1A exhibit a higher circularity and a lower aspect ratio compared to mCherry control or RUFY3-mCherry expressing cells, although the cell area remained unaffected (Figure S5L-N). In addition, as previously described [63] KIF1A redirected lysosomes to FA sites (Figure 5F-G, Figure S5A), which might suggest increased FA remodeling [25,59]. Instead, overexpression of mCherry-KIF1A resulted in higher numbers and larger size of FAs compared to both the mCherry control and RUFY3-mCherry (Figure 5F and H). Thus, KIF1A increased LAMP1 at FAs and elevated FA number/area while reducing protrusions, consistent with a more adhesive, less protrusive state.

### Forced peripheral dispersion of lysosomes impacts motility and membrane protrusive activity

We next performed live cell imaging experiments to investigate the effects of the manipulation of lysosomal positioning on protrusion formation, cell motility, and on the subcellular localization of GFP-PLEKHG3 (Supp Movies S6-11, Figure 6A). Cells expressing mCherry-KIF1A were rounder than mCherry control and RUFY3-mCherry expressing cells, and were much less active than the other cells, particularly the RUFY3-mCherry expressing cells. Morphometric analysis performed over a time span of 128 min revealed that KIF1A and RUFY3 slightly increased cell size in the GFP cell line; the most evident increase in size, however, was caused by GFP-PLEKHG3 itself, particularly when co-expressed with RUFY3 (Figure 6B). GFP-PLEKHG3 also promoted circularity, even more so upon co-expression of mCherry-KIF1A, confirming the experiments in fixed cells (Figure 5A + Figure S5M). Accordingly, analysis of the aspect ratio of the cells averaged over time showed that mCherry-KIF1A caused a decrease, while RUFY3 caused a slight increase in this parameter, which was not impacted by GFP-PLEKHG3 expression (Figure 6D).

**Figure 6:**
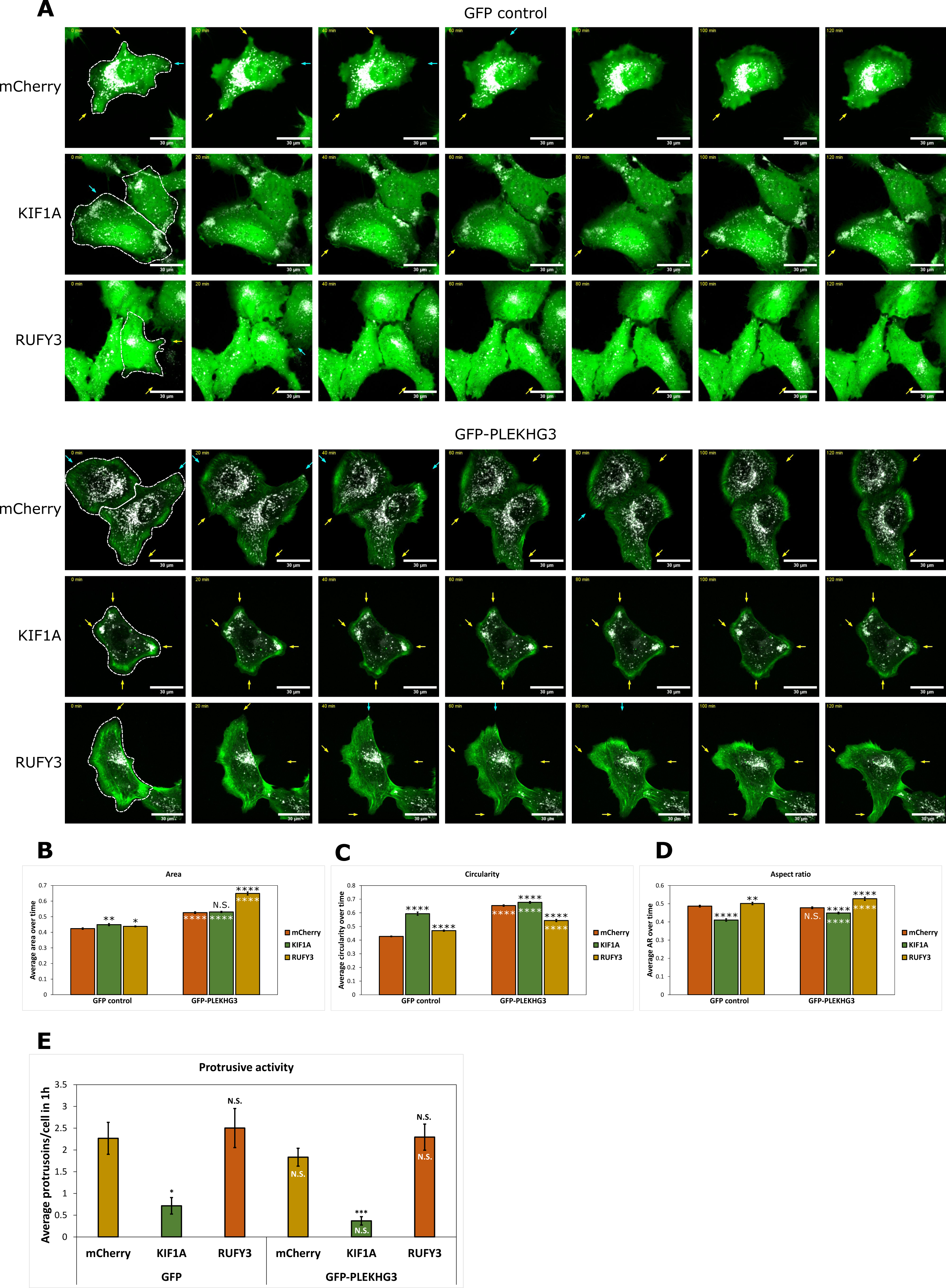
Peripheral clustering of lysosomes inhibits protrusion formation. **A)** Stills from live cell imaging (Movies S6-11). Cells stably expressing GFP or GFP-PLEKHG3 were transfected with the indicated mCherry constructs. Successfully transfected cells are contoured by a white broken line in the first still image. Yellow arrows = forming protrusions; blue arrows = retracting protrusions. Stills were generated over a period of 2 hrs. Scale bar = 30 µm. **B-D)** Morphometric analysis of cells over time. Cell shape analysis was performed with Fijis shape descriptors (see materials and methods) per cell every four minutes. Bar plots represent the average over all time points to represent changes in shape over time. **E)** Quantification of protrusions formed and retracted over time in cells from A. Values indicate average number of protrusions formed in a timespan of one hour from a total of ≥ 19 cells per condition. Error bars = SEM. PLE3 = PLEKHG3. In **B-E**, black asterisks denote p values according to student’s t-test, comparing the effect of KIF1A or RUFY3 expression with the effect of mCherry. White asterisks in the bars indicate the p values according to student’s t-test comparing GFP-PLEKHG3 and GFP expressing cells.

Analysis of the number of emerging protrusions and retracting membranes, averaged over a one-hour timespan, revealed a significant decrease in protrusion formation upon overexpression of mCherry-KIF1A (Figure 6E + Figure S6A-B). Reduced protrusive activity correlated with less dynamic LifeAct-positive [64] F-actin structures, as shown in GFP and GFP-PLEKHG3 cells, co-transfected with BFP-LifeAct, and mCherry control or mCherry-KIF1A (Supp Movies S12-15, Figure S6A-C + Supp Movies S16-17, Figure S6D). This effect was further exacerbated in cells expressing GFP-PLEKHG3 (Figure 6E + Figure S6B), thereby confirming the results presented in Figure 5E. GFP-PLEKHG3 colocalized with the BFP-LifeAct structures in the presence or absence of mCherry-KIF1A (Figure S6C-D, Supp Movies S16-17). The decrease in protrusion formation and F-actin dynamics induced by mCherry-KIF1A was also observed in PLEKHG3 KO cells, indicating that the effect of KIF1A was PLEKHG3-independent under these conditions (Figure S6E-G).

## Discussion

In this study, we identify PLEKHG3 as a vicinal protein of L3 by proximity-dependent labeling and show that GFP-PLEKHG3 accumulates at FAs, where it colocalizes with peripheral lysosomes. We demonstrate that peripheral clustering of lysosomes concentrates PLEKHG3 at the plasma membrane, while protrusion formation and cell motility decrease under these conditions. The suppression of protrusions occurs in both wild-type and PLEKHG3 knockout cells, indicating PLEKHG3 is not required for this effect in our system.

The proximity labeling screen correctly identified most of the proteins known to associate with the LAMTOR complex (Figure 1D). However, both lysosomal baits, L3 and TMEM192, failed to biotinylate LAMP1, a very abundant lysosomal transmembrane protein. This is in line with a BioID screen performed by Go et al. in HEK293 cells [49], which showed that LAMP1 was not biotinylated by a LAMTOR1 bait and vice versa. This supports the view that the proteins at the lysosome’s membrane are spatially compartmentalized into distinct subdomains.

Similarly, TMEM192 missed several of the proteins labeled by L3, including the prominent biotinylated PLEKHG3. This could be explained by the fact that TMEM192 features only a small cytosolic tail and thus doesn’t reach as far into the cytoplasm as L3, which is more distal from the membrane [18,65]. Be that as it may, labeling of PLEKHG3 by L3 did not reflect a stable interaction (Figure 2A, C-D), and L3 KO in turn did not have an influence on PLEKHG3 distribution (Figure S3E-L, Movie S1-4). Therefore, PLEKHG3 is a vicinal protein of L3 rather than a stable interactor.

### PLEKHG3 colocalizes with lysosomes at FA sites

PLEKHG3 has previously been shown to localize to both the leading and the trailing edge of migrating cells, where remodeling of the actin cytoskeleton takes place, and to associate with freshly polymerized actin via its actin-binding domain [53]. Our experiments confirm this localization of GFP-PLEKHG3 to cortical actin, but in addition show that PLEKHG3 is specifically enriched at FA sites in HeLa cells (Figure 3A and Fig 4B-D). During protrusion formation, new FAs are formed by extracellular matrix/integrin interaction. These nascent adhesions connect to actin fibers and further mature to FAs (reviewed in [66]). The connection with growing actin fibers may be the basis of the accumulation of PLEKHG3 around FAs.

In line with published evidence [63], we find an enrichment of peripheral lysosomes at FA sites (Figure 4B-E), where they fulfil a number of functions, including the control of FA dynamics by IQGAP release from FA complexes [59] or by dissolution of extracellular matrix/Integrin complexes [37,67,68].

The observation that PLEKHG3 and LAMP1-positive vesicles were in close proximity to one another at FA and protrusion sites (Figure 4B-E, Figure 3C-F) led to the hypothesis that PLEKHG3 biotinylation by L3 was the result of dynamic encounters at sites of protrusion formation and prompted us to ask whether association with the lysosomes or lysosomal trafficking had any impact on PLEKHG3 distribution. Indeed, the effect was profound (Figure 5A-D). Driving lysosome outward transport by overexpression of KIF1A promoted the redistribution of both GFP-PLEKHG3 and of endogenous PLEKHG3 to the periphery of the cell. Whether the lysosomes are directly responsible for the transport of PLEKHG3 to the membrane is difficult to determine, since both endogenous and exogenous PLEKHG3 accumulated at protrusions which frequently harbored lysosomes but could also be found at sites that contained very few to no lysosomes (Figure 3C-F). Promoting lysosome translocation to the periphery did not increase protrusive activity; instead, protrusions and actin dynamics decreased while FA number/area increased, suggesting that altered adhesion dynamics dominate protrusion output. Clustering of lysosomes around the nucleus by RUFY3 had the opposite effect both on PLEKHG3 distribution and cell morphology/motility (Figure 5E + Figure S5L-N).

The negative effect of KIF1A and PLEKHG3 overexpression on protrusion activity and cell motility was unexpected in two respects: firstly, PLEKHG3 overexpression has been shown to promote protrusion activity by binding to newly synthesized F-actin filaments [53], a result we confirmed by GFP-PLEKHG3 and BFP-LifeAct colocalization analysis, even upon KIF1A overexpression (Figure S6C-D); and secondly, centrifugal trafficking of lysosomes towards the periphery has been shown to contribute to protrusion activity, spreading and motility [15,39,69]. Lysosomes can contribute to protrusion formation e.g. by delivering membranes and actin polymerization-promoting factors to the newly formed structures (reviewed in [70–73]. Integrins, for instance, have been shown to travel via the late endosome/lysosome route to the periphery where they aid to establish new FAs and thereby promote protrusion formation [74,75]. In line with the latter, we identified an increased number and size of FAs in cells overexpressing KIF1A (Figure 5F + H), which might contribute to a more static cellular behavior. It is likely that upsetting the balance between anterograde and retrograde transport of lysosomes within this cellular context prevents the removal and degradation of membrane components involved in FA dynamics (e.g. integrins) and/or in the actin remodeling necessary for protrusion formation and motility. As an example, it has recently been reported that lysosomes remove the GDP-bound, inactive form of the PLEKHG3 downstream effector Rac1 from the vicinity of the PM in order to increase the concentration of GTP-bound, active Rac1 at the leading edge, thus promoting cell migration [43].

In this situation of impasse, PLEKHG3 might be recruited to the PM, but this alone might not be sufficient to promote protrusion activity and cell motility.

Taken together, our results identify PLEKHG3 as a new L3 vicinal protein that encounters lysosomes at FAs, revealing a previously undiscovered interplay between lysosomal trafficking and PLEKHG3 distribution/protrusion-promoting function. The data highlight the unique potential of PDL-based approaches to discover interactions between molecules, or as shown here between molecules and intracellular structures, that are physiologically relevant but limited in space and time.

## Supporting information

Graphical abstract

## Acknowledgments

We are grateful M.E.G. de Araujo and T. Stasyk for helpful discussions, advice during the project and the gift of the HeLa cell line used in this study as well as a LAMTOR3 plasmid. We thank A. Ballabio and D. Medina for their support at TIGEM. We are grateful to C. Guardia who kindly provided us with a KIF1A plasmid, the MPL FACS facility and the MPL BioOptics facility, especially I. Fischer and T. Peterbauer for assistance with imaging and macro writing. We thank the MPL Mass Spectrometry Facility which performed the proteomics analysis with assistance of N. Hartl and D. Anrather using the VBCF instrument pool. We thank S. Martens for advice during the project.

This work was supported by the Austrian Science Fund W-1220 “Signaling mechanisms in cellular homeostasis” (to M. Baccarini) and by funds of the University of Vienna (to M. Baccarini). G. Vucak is a recipient of a DOC-fellowship of the Austrian Academy of Sciences at the Max Perutz Labs. L.A. Huber is funded by the Austrian Science Fund P-32608.

## Author contributions

R. Ettelt performed most of the experiments. G. Vucak contributed the L3-T-V5 screen data and immunoblots. A.S. Sandru performed live cell imaging experiments. S. Didusch performed the bioinformatic analysis of MS data and helped in microscopic data analysis. B. Riemelmoser performed selected cell biology and biochemistry experiments. Karin Ehrenreiter helped with the generation of stable cell lines and selected experiments. M. Hartl performed the mass spectrometry. M. Baccarini, R. Ettelt and L. A. Huber wrote the manuscript. The whole project was supervised by M. Baccarini.

## Abbreviations

FA: (Focal adhesion)
GEF: (Guanine-nucleotide exchange factor)
KIF1A: (Kinesin family member 1A)
LAMTOR: (Late endosomal/lysosomal adaptor and MAPK and mTORC1 activator)
L3: (LAMTOR3)
PLEKHG3: (Pleckstrin Homology and RhoGEF Domain Containing G3)
PM: (Plasma membrane)
RUFY3: (RUN and FYVE domain-containing protein 3)

## Online supplemental material

Figure S1 shows validation of TurboID baits and bioinformatic analysis. Figure S2 shows validation of PLEKHG3 reagents and PLEKHG3 KO. Figure S3 shows effect of L3 KO on localization and function of GFP-PLEKHG3. Figure S4 shows that PLEKHG3 KO does not influence lysosomal distribution or cell morphometry. Figure S5 shows the effect of lysosomal distribution on PLEKHG3 and cell morphology. Figure S6 shows that PLEKHG3 is dispensable for protrusive activity. Movie S1-4 show L3 CTRL and KO cells expressing GFP or GFP-PLEKHG3. Movie S5 shows lysosomes moving into forming protrusions. Movies S6-11 show effect of changed lysosomal localization on protrusion formation and cell shape. Movies S12-15 show colocalization of GFP/GFP-PLEKHG3 with LifeAct-positive F-actin structures upon lysosomal relocation. Movies S16-17 show selected cells from experiments presented in Movies S14-15 portraited as colocalization heatmaps. Movies S18-21 show the effect of PLEKHG3 KO on GFP/GFP-PLEKHG3 colocalization upon lysosomal relocation. Table S1 contains all identified proteins, spectral counts, LFQ intensities, and differential expression analysis results for L3-T-V5 as displayed in Figure S1D. Table S2 contains all identified proteins, spectral counts, LFQ intensities, and differential expression analysis results for V5-T-L3 as displayed in Figure S1I-K.

