## Supplementary information and Figures for "Peripheral lysosomes recruit PLEKHG3 to focal adhesions and restrain protrusion dynamics"

**Figure S1:** Validation of baits used in TurboID screens and bioinformatic analysis

**Figure S2:** Validation of PLEKHG3 reagents and PLEKHG3 KO

**Figure S3:** L3 is dispensable for the colocalization of PLEKHG3 with LAMP1-positive structures and for the localization of PLEKHG3 to FAs

**Figure S4:** PLEKHG3 KO does not influence lysosomal positioning or cell morphometrics

**Figure S5:** Effect of lysosomal positioning on PLEKHG3 and cell morphology

**Figure S6:** PLEKHG3 localizes to F-actin independently of lysosomal transport and is dispensable for protrusive activity.

**Movie S1:** stable HeLa L3 CTRL GFP + LysoTracker Deep Red
**Movie S2:** stable HeLa L3 KO GFP + LysoTracker Deep Red
**Movie S3:** stable HeLa L3 CTRL GFP-PLEKHG3 + LysoTracker Deep Red
**Movie S4:** stable HeLa L3 KO GFP-PLEKHG3 + LysoTracker Deep Red

**Movie S5:** HeLa transiently transfected with GFP-PLEKHG3 + LysoTracker Red DND-99

**Movie S6:** HeLa parental GFP + mCherry control + LysoTracker Deep Red
**Movie S7:** HeLa parental GFP + mCherry -KIF1A + LysoTracker Deep Red
**Movie S8:** HeLa parental GFP + RUFY3-mCherry + LysoTracker Deep Red
**Movie S9:** HeLa parental GFP-PLEKHG3 + mCherry control + LysoTracker Deep Red
**Movie S10:** HeLa parental GFP-PLEKHG3 + mCherry -KIF1A + LysoTracker Deep Red
**Movie S11:** HeLa parental GFP-PLEKHG3 + RUFY3-mCherry + LysoTracker Deep Red

**Movie S12:** HeLa parental GFP + mCherry control + LifeAct
**Movie S13:** HeLa parental GFP + mCherry-KIF1A + LifeAct
**Movie S14:** HeLa parental GFP-PLEKHG3 + mCherry control + LifeAct
**Movie S15:** HeLa parental GFP-PLEKHG3 + mCherry-KIF1A + LifeAct

**Movie S16:** Coloc-heatmap of HeLa parental GFP-PLEKHG3 + mCherry
**Movie S17:** Coloc-heatmap of HeLa parental GFP-PLEKHG3 + mCherry-KIF1A

**Movie S18:** PLEKHG3 WT HeLa + mCherry control + LifeAct + LysoTracker

**Movie S19:** PLEKHG3 WT HeLa + mCherry-KIF1A + LifeAct + LysoTracker
**Movie S20:** PLEKHG3 KO HeLa + mCherry control + LifeAct + LysoTracker
**Movie S21:** PLEKHG3 KO HeLa + mCherry-KIF1A + LifeAct + LysoTracker

**Table S1:** Raw data and differential expression analysis of L3-T-V5
**Table S2:** Raw data and differential expression analysis of V5-T-L3

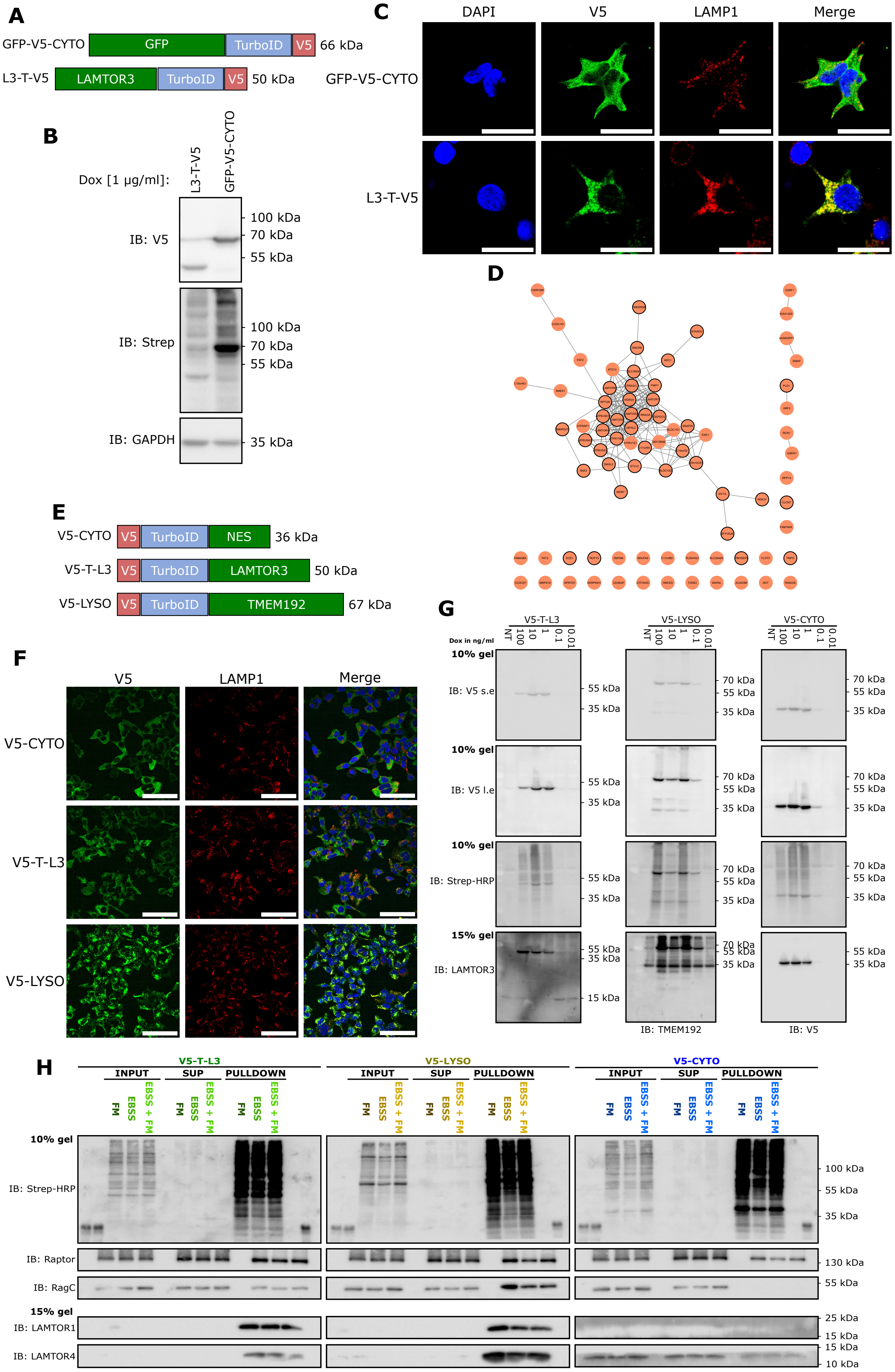
A
GFP-V5-CYTO
GFP
TurboID
V5
66 kDa
L3-T-V5
LAMTOR3
TurboID
V5
50 kDa
B
Dox [1 µg/ml]:
L3-T-V5
GFP-V5-CYTO
IB: V5
100 kDa
70 kDa
55 kDa
IB: Strep
100 kDa
70 kDa
55 kDa
IB: GAPDH
35 kDa
C
DAPI
V5
LAMP1
Merge
GFP-V5-CYTO
L3-T-V5
D
E
V5-CYTO
V5
TurboID
NES
36 kDa
V5-T-L3
V5
TurboID
LAMTOR3
50 kDa
V5-LYSO
V5
TurboID
TMEM192
67 kDa
F
V5
LAMP1
Merge
V5-CYTO
V5-T-L3
V5-LYSO
G
V5-T-L3
V5-LYSO
V5-CYTO
Dox in ng/ml
NT 100 10 1 0.1 0.01
10% gel
IB: V5 s.e
10% gel
IB: V5 l.e
10% gel
IB: Strep-HRP
15% gel
IB: LAMTOR3
IB: TMEM192
IB: V5
70 kDa
55 kDa
35 kDa
15 kDa
H
V5-T-L3
V5-LYSO
V5-CYTO
INPUT
SUP
PULLDOWN
FM
EBSS
EBSS + FM
10% gel
IB: Strep-HRP
IB: Raptor
IB: RagC
15% gel
IB: LAMTOR1
IB: LAMTOR4
100 kDa
55 kDa
35 kDa
130 kDa
55 kDa
25 kDa
15 kDa
15 kDa
10 kDa

**I**

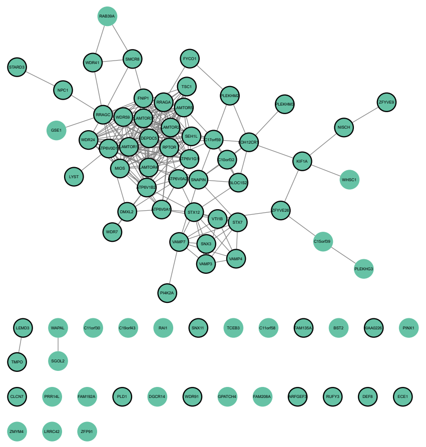

**J**

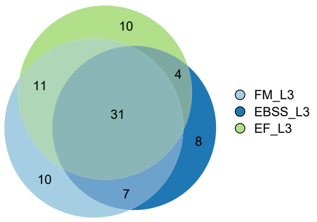


**K**

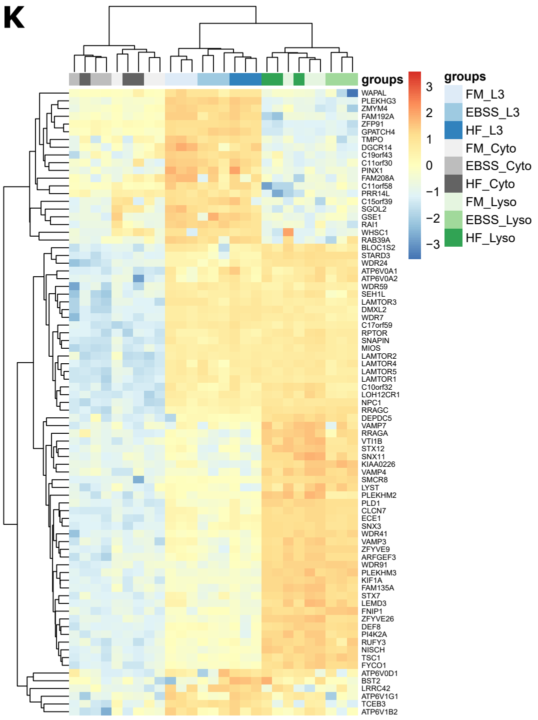


***Figure S1:*** **Validation of baits used in TurboID screen and bioinformatic analysis. A)** L3 fusion proteins used in the first screen (L3-T-V5). GFP-tagged TurboID served as a cytosolic control (GFP-V5-CYTO). **B)** Immunoblot of HEK293T cells stably expressing L3-T-V5 and GFP-V5-CYTO. Expression of GFP-V5-CYTO is slightly higher compared to L3. **C)** Confocal images of HEK293T cell showing L3-T-V5 bait overlaps with LAMP1. HEK293T cells grown under steady-state conditions were immunostained with V5 and LAMP1 antibodies. Scale bar = 30 µm. **D)** STRING-network representation of hits identified by comparing L3-T-V5 to GFP-V5-CYTO (log2FC > 1.49 and adj. p-value ≤ 0.05). Nodes outlined in black indicate hits detected in the second screen that were significantly enriched in the V5-LYSO over the V5-CYTO bait (log2FC > 1.49 and adj. p-value ≤ 0.05). **E)** V5-T-L3 fusion protein, cytosolic control (V5-CYTO) and lysosomal signpost (V5-LYSO) used in the second screen. **F)** Confocal images of bait localization in T-REx cells. Cells grown under steady-state conditions were immunostained for V5 and LAMP1 to visualize the overlap of the bait with lysosomes. Expression of constructs was induced with 1 ng/mL doxycycline. Scale bar = 100 µm. **G)** Doxycycline titration of bait to match endogenous expression levels. T-REx cells were cultured in full medium and treated with the indicated concentrations of doxycycline for 24 hours. Prior to cell lysis, biotin was added to the medium. Samples were analyzed by immunostaining. Expression of fusion protein is indicated by V5 staining and the ligase activity by Streptavidin probing. **H)** Benchmarking final conditions for the MS experiments. Cells were exposed to the indicated conditions (EBSS = 30 minutes and EBSS + FM = 30 minutes + 10 minutes) and lysed 15 minutes after the addition of biotin to the medium. Biotinylated proteins were subjected to immunoblot analysis. Known interactors and proteins of interest were used as benchmarks for PDL efficiency. **I)** STRING-network representation of hits identified by V5-T-L3 compared to V5-CYTO (log2FC > 1.49 and adj. p-value ≤ 0.05) in cells kept under different culture conditions. Proteins outlined in black were also significantly enriched in the V5-LYSO over the V5-CYTO bait (log2FC > 1.49 and adj. p-value ≤ 0.05). **J)** Venn-diagram of preys biotinylated by V5-T-L3 and significantly enriched over V5-CYTO in all three culture conditions (log2FC > 1.49 and adj. p-value ≤ 0.05). FM = full medium, EBSS = growth factor + amino acid starvation, HF = EBSS + 10min FM. **K)** Heatmap representation of hits identified in all three media by comparing V5-T-L3 with V5-CYTO (log2FC > 1.49 and adj. p-value ≤ 0.05). All nuclear hits excluded.

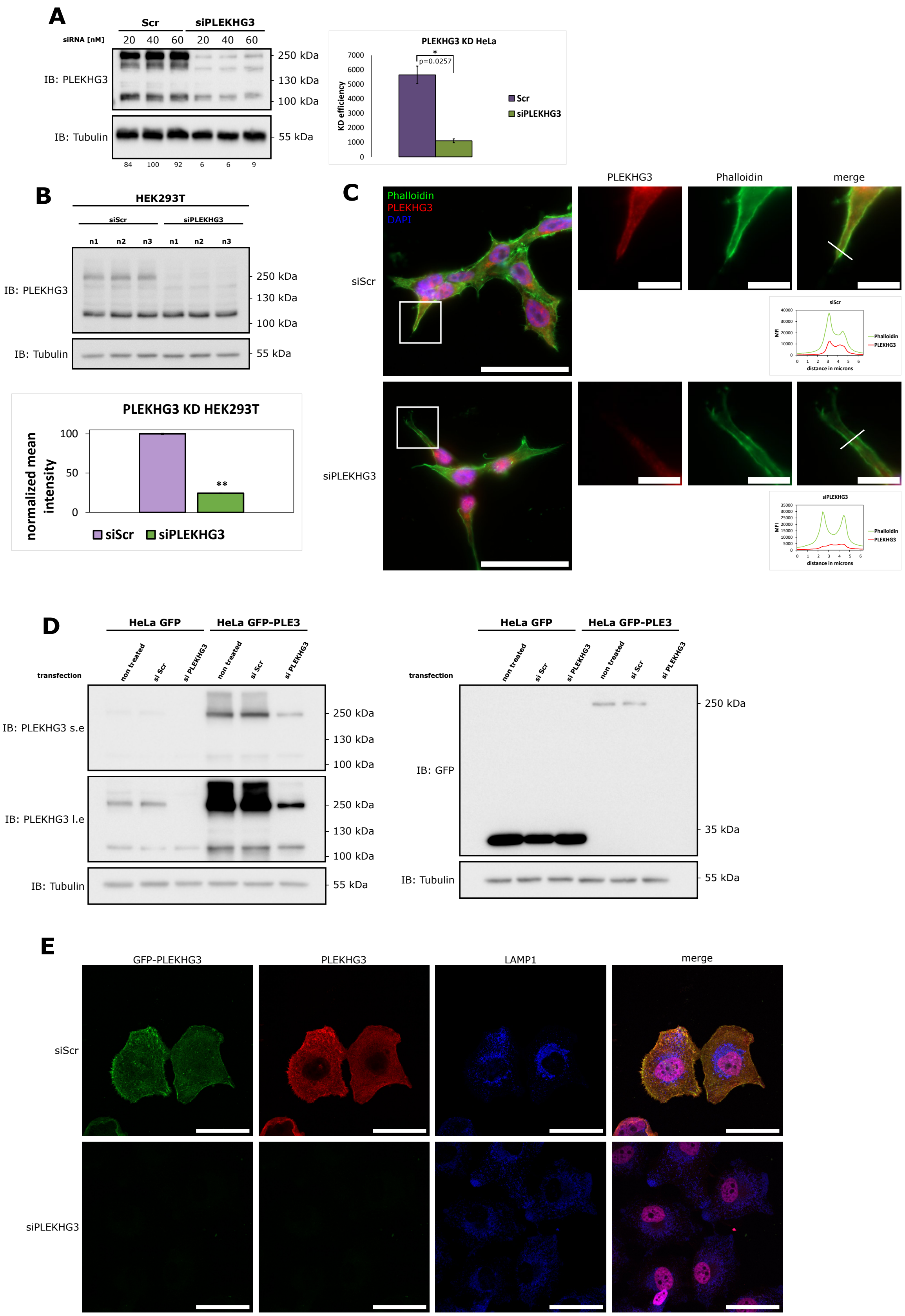
A
Scr
siPLEKHG3
siRNA [nM]
20 40 60 20 40 60
IB: PLEKHG3
IB: Tubulin
250 kDa
130 kDa
100 kDa
55 kDa
84 100 92 6 6 9
PLEKHG3 KD HeLa
KD efficiency
p=0.0257
Scr
siPLEKHG3
B
HEK293T
siScr
siPLEKHG3
n1 n2 n3 n1 n2 n3
IB: PLEKHG3
IB: Tubulin
PLEKHG3 KD HEK293T
normalized mean intensity
siScr
siPLEKHG3
C
Phalloidin
PLEKHG3
DAPI
merge
siScr
siPLEKHG3
distance in microns
D
HeLa GFP
HeLa GFP-PLE3
transfection
non treated
si Scr
si PLEKHG3
IB: PLEKHG3 s.e
IB: PLEKHG3 l.e
IB: Tubulin
IB: GFP
35 kDa
E
GFP-PLEKHG3
PLEKHG3
LAMP1
merge
siScr
siPLEKHG3

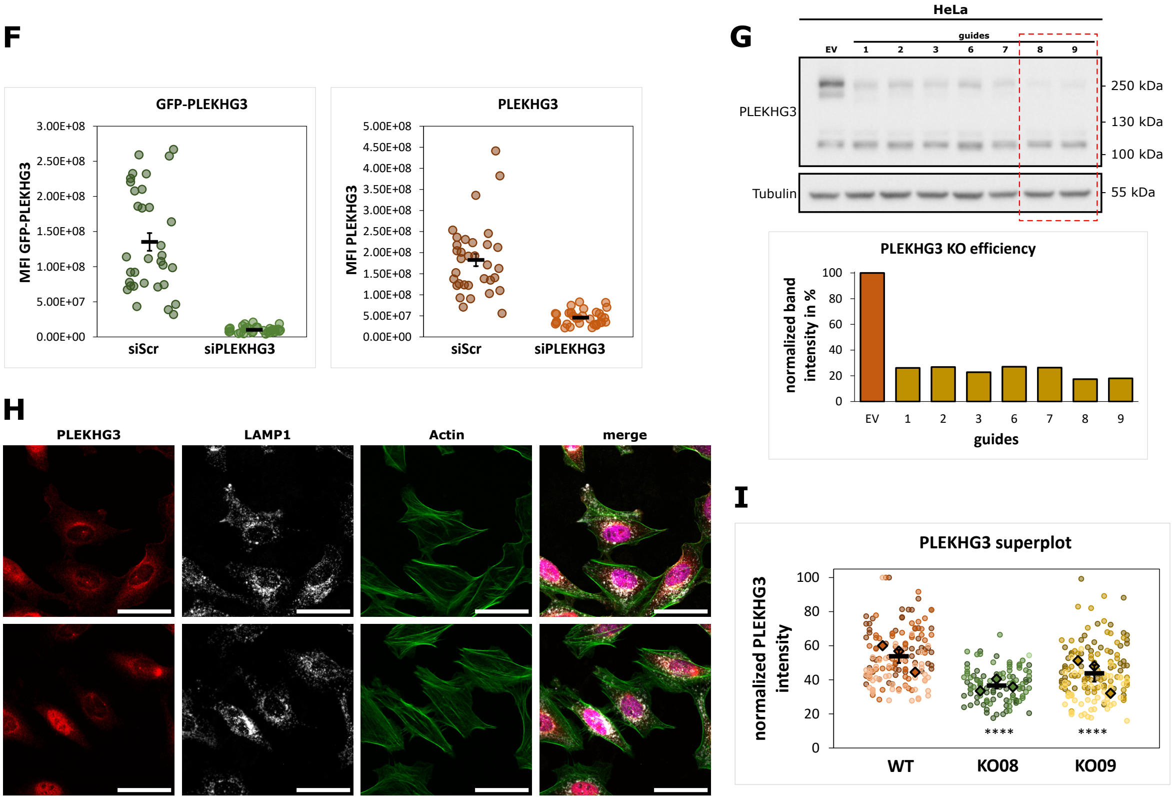


***Figure S2:*** **Validation of PLEKHG3 reagents. A)** Immunoblot of HeLa samples treated with the indicated siRNA concentrations probed with antibodies against PLEKHG3 and tubulin as loading control. Numbers below blot indicate KD efficiency in percent. KD efficiency was calculated by drop of PLEKHG3 band intensity (uppermost band corrected to the loading control) normalized to max PLEKHG3 band intensity. Bar plot on the right represents quantification of PLEKHG3 KD efficiency from three biological replicates. Error bars = SEM, n=3. * = p values according to Student's t-test. **B)** Western blot of HEK293T cells showing downregulation of PLEKHG3 expression upon siPLEKHG3 treatment compared to siScr. Bar plot shows quantification of PLEKHG3 bands from immunoblot above. Error bars = SEM, n=3. * = p values according to Student's t-test. **C)** Immunofluorescence images of HEK293T cells. siPLEKHG3 shows a drop in PLEKHG3 intensity in the periphery of the cell and less colocalization with Phalloidin. Scale bar = 50 µm. Line plots show intensity profiles of Phalloidin (green) and PLEKHG3 (red) along the white lines in the merged inset images. Scale bar = 10 µm. **D)** Immunoblot of HeLa cells stably expressing GFP or GFP-PLEKHG3 treated with indicated siRNAs. The drop in endogenous PLEKHG3 as well as exogenously expressed GFP-PLEKHG3 signal could be observed by both, PLEKHG3 and GFP antibody staining. Tubulin was used as a loading control. **E)** Confocal immunofluorescence images of siScr or siPLEKHG3 treated HeLa cells stably expressing GFP-PLEKHG3. LAMP1 antibody was used for lysosomal staining, PLEKHG3 antibody and GFP fluorescence for GFP-PLEKHG3 localization and DAPI to identify the nucleus. Scale bar = 50µm. **F)** Superplot quantification of E. The mean fluorescence intensity of the GFP and PLEKHG3 signal/cell is represented by single dots (n=33). **G)** Immunoblot and quantification of HeLa PLEKHG3 KO cells represents the degree of PLEKHG3 depletion achieved using different guides compared to WT cells transfected with empty vector (EV). The most potent guides (8-9) are boxed in red. **H)** Immunofluorescence images of WT and PLEKHG3 KO8 cells reveal an overall drop in PLEKHG3 intensity and the specific loss of PLEKHG3 signal at the periphery of the cells. **I)** Quantification of PLEKHG3 intensity as displayed in H for two KO cell lines compared to WT cells. Dots represent individual data points of each of the three-color coded replicates; diamonds represent the mean of each replicate; black bars represent the mean ± the SEM of three biological replicates; * = p values according to two-way-ANOVA.

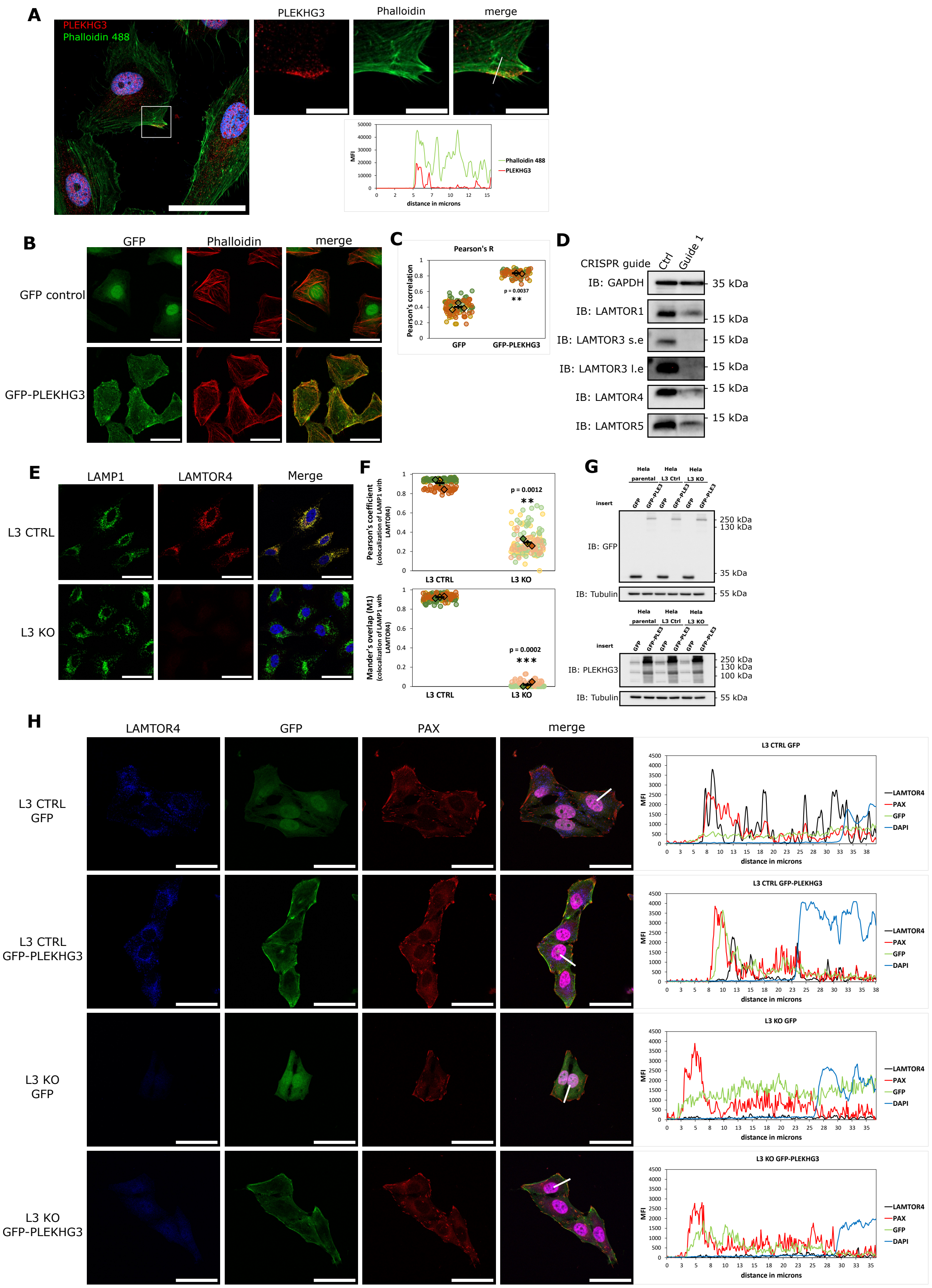
A
PLEKHG3
Phalloidin 488
PLEKHG3
Phalloidin
merge
MFI
distance in microns
Phalloidin 488
PLEKHG3
B
GFP
Phalloidin
merge
GFP control
GFP-PLEKHG3
C
Pearson's R
Pearson's correlation
p = 0.0037
GFP
GFP-PLEKHG3
D
CRISPR guide
Ctrl
Guide 1
IB: GAPDH
35 kDa
IB: LAMTOR1
15 kDa
IB: LAMTOR3 s.e
15 kDa
IB: LAMTOR3 l.e
15 kDa
IB: LAMTOR4
15 kDa
IB: LAMTOR5
15 kDa
E
LAMP1
LAMTOR4
Merge
L3 CTRL
L3 KO
F
Pearson's coefficient (colocalization of LAMP1 with LAMTOR4)
p = 0.0012
L3 CTRL
L3 KO
Mander's overlap (M1) (colocalization of LAMP1 with LAMTOR4)
p = 0.0002
L3 CTRL
L3 KO
G
Hela parental
Hela L3 Ctrl
Hela L3 KO
insert
GFP
GFP-PLE3
IB: GFP
250 kDa
130 kDa
35 kDa
IB: Tubulin
55 kDa
IB: PLEKHG3
250 kDa
130 kDa
100 kDa
IB: Tubulin
55 kDa
H
LAMTOR4
GFP
PAX
merge
L3 CTRL GFP
L3 CTRL GFP-PLEKHG3
L3 KO GFP
L3 KO GFP-PLEKHG3
MFI
distance in microns
LAMTOR4
PAX
GFP
DAPI

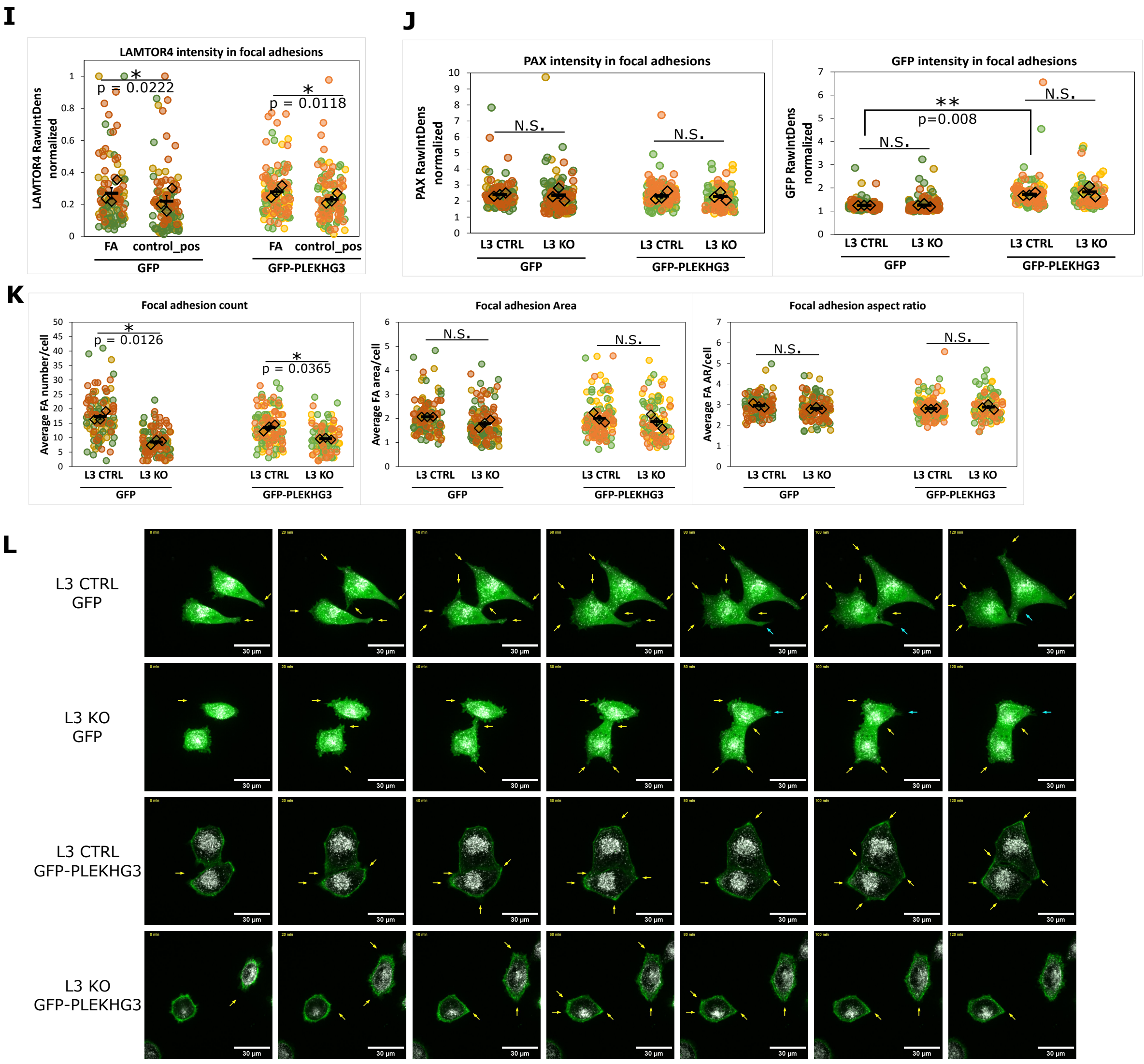


***Figure S3:*** **L3 is dispensable for the colocalization of PLEKHG3 with LAMP1-positive structures and for the localization of PLEKHG3 to FAs. A)** HeLa cells stained with PLEKHG3 (red) and Phalloidin (green). The nucleus is indicated by DAPI staining (blue). Scale bar = 50 µm. Insets on the right as indicated by white box in image on the left. Scale bar = 10 µm. Line plot corresponds to white line in merged inset. **B)** Confocal immunofluorescence images of HeLa cells stably expressing GFP or GFP-PLEKHG3. Cells were stained with Phalloidin to visualize the actin cytoskeleton. Scale bar = 50 µm. **C)** Colocalization analysis of GFP-PLEKHG3 and Phalloidin from experiments similar to the one shown in A confirms the localization of GFP-PLEKHG3 to the actin cytoskeleton. Pearson's correlation was calculated by Fiji's JACoP plugin. **D)** Immunoblot of L3 CTRL and KO cells obtained using two different RNA guides. Both show loss of L3 and reduction of LAMTOR expression (LAMTOR1, 4 and 5). Guide 1 showed the strongest effect and was therefore used in the following experiments. GAPDH served as a loading control. **E)** Confocal immunofluorescence images of L3 CTRL and L3 KO HeLa cells stained with LAMP1 (green) and LAMTOR4 (red) antibodies. Upon deletion of L3, LAMTOR4 dissociates from lysosomes and its expression is reduced. Scale bar = 50 µm. **F)** Quantification of LAMP1 and LAMTOR4 colocalization from experiments similar to the one represented in D by both, Pearson's correlation (upper panel) and Mander's coefficient (lower panel). **G)** Immunoblot of parental, L3 CTRL and L3 KO HeLa cells lines stably expressing GFP or GFP-PLEKHG3. Membranes were probed with PLEKHG3 or GFP antibodies. Tubulin was used as a loading control. **H)** Confocal images of L3 CTRL or L3 KO HeLa cells stably expressing GFP or GFP-PLEKHG3 (green). Cells were stained with LAMTOR4 antibodies (white) to label lysosomes and at the same time show the effects of the L3 KO, and with Paxillin antibodies to label FAs (red). Scale bar = 50 µm. The line plots on the right show the intensity of the respective fluorophores along the white lines drawn on the merged images. **I)** Colocalization of LAMTOR4 with FAs in L3 CTRL cells expressing GFP or GFP-PLEKHG3. Colocalization with FAs was measured by comparing LAMTOR4 or GFP/GFP-PLEKHG3 content in Paxillin-positive regions and in adjacent, paxillin-negative regions of equal area and shape. **J)** L3 KO does not impact Paxillin or GFP/GFP-PLEKHG3 content in FAs. Colocalization with FAs was measured as in H. **K)** Quantification of FA morphometrics of L3 CTRL and L3 KO cells shown in Figure S3D. Except for the number of FAs there was no significant difference detectable between L3 CTRL and KO cells in both GFP and GFP-PLEKHG3 expressing cells. **L)** Stills from movies S1-4. L3 CTRL or L3 KO cells expressing GFP or GFP-PLEKHG3 were stained with LysoTracker Deep Red and imaged for 3 hours. Yellow arrows show forming and blue arrows retracting protrusions. Scale bar = 30 µm. In the superplots in **C, F, I-K**, dots represent individual data points of each of the three-color coded replicates; diamonds represent the mean of each replicate; black bars represent the mean ± the SEM of three biological replicates; * = p values according to Student's t-test.

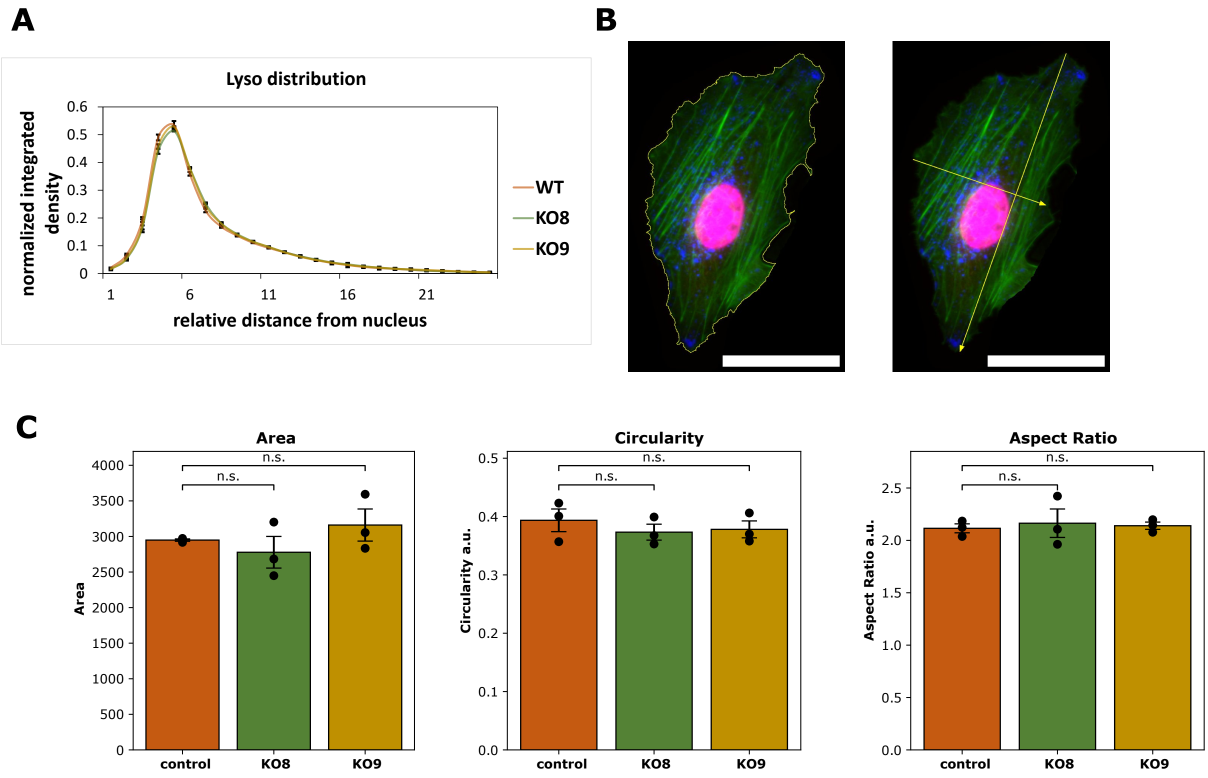


**Figure S4: PLEKHG3 KO does not influence lysosomal distribution or cell morphometry: A)** Quantification of lysosomal distribution in WT compared to two KO cell lines. N ≥ 50 cells in three biological replicates. **B)** Schematic representation of analysis of cell shape descriptors as referred to in C). Left picture shows the calculated outline in yellow based on which the cell area and circularity are calculated. Right picture shows the minor and major cell axis which, calculated as fraction, result in the aspect ratio of the cell. Scale bar = 50 µm. **C)** Quantification of cell morphometric parameters Area, Circularity and Aspect ratio. N ≥ 50 cells in three biological replicates. Black dots represent mean of each biological replicate. Statistical analysis according to Student's t-test. Error bars = SEM.

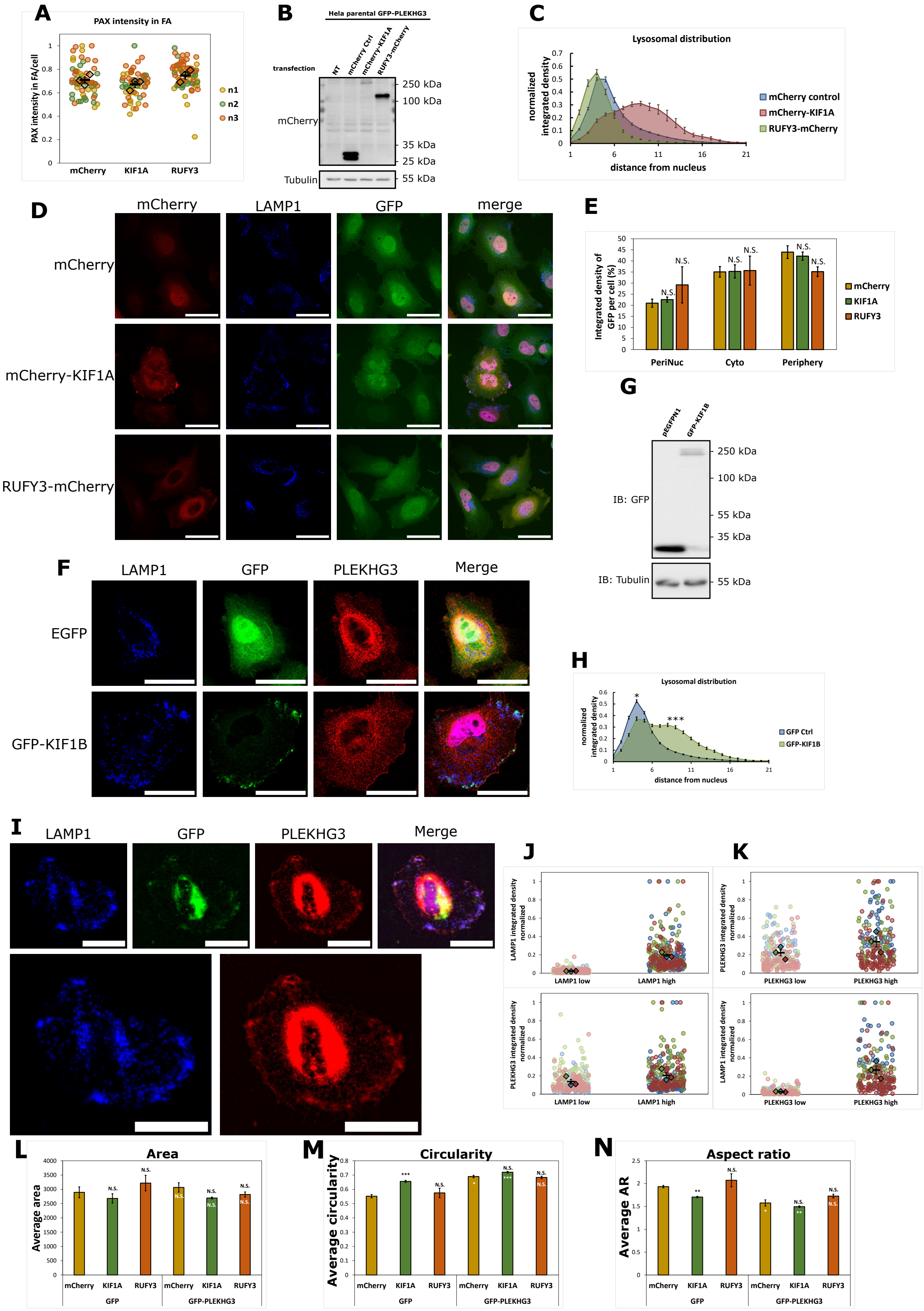
A
PAX intensity in FA
PAX intensity in FA/cell
mCherry
KIF1A
RUFY3
n1
n2
n3
B
Hela parental GFP-PLEKHG3
transfection
NT
mCherry Ctrl
mCherry-KIF1A
RUFY3-mCherry
mCherry
Tubulin
250 kDa
100 kDa
35 kDa
25 kDa
55 kDa
C
Lysosomal distribution
normalized integrated density
distance from nucleus
mCherry control
mCherry-KIF1A
RUFY3-mCherry
D
mCherry
LAMP1
GFP
merge
mCherry
mCherry-KIF1A
RUFY3-mCherry
E
Integrated density of GFP per cell (%)
N.S.
PeriNuc
Cyto
Periphery
mCherry
KIF1A
RUFY3
G
pEGFPN1
GFP-KIF1B
IB: GFP
IB: Tubulin
250 kDa
100 kDa
55 kDa
35 kDa
55 kDa
F
LAMP1
GFP
PLEKHG3
Merge
EGFP
GFP-KIF1B
H
Lysosomal distribution
normalized integrated density
distance from nucleus
GFP Ctrl
GFP-KIF1B
I
LAMP1
GFP
PLEKHG3
Merge
J
K
LAMP1 integrated density normalized
LAMP1 low
LAMP1 high
PLEKHG3 integrated density normalized
PLEKHG3 low
PLEKHG3 high
L
Area
Average area
M
Circularity
Average circularity
N
Aspect ratio
Average AR
mCherry
KIF1A
RUFY3
GFP
GFP-PLEKHG3

**Figure S5: Effect of lysosomal positioning on PLEKHG3 and cell morphology. A)** Control analysis for Figure 5G. Superplot represents average Paxillin intensity per cell in FA structures for three biologically independent experiments. **B)** Immunoblot analysis of mCherry expression. **C)** Quantification of lysosomal distribution in cells overexpressing the indicated mCherry constructs (Figure 5A). KIF1A overexpression concentrated lysosomes in the periphery, RUFY3 overexpression in the perinuclear region of the cell. Values represent mean of > 50 cells from three biological replicates. Error bars = SEM. **D)** Confocal immunofluorescence images of HeLa cells stably expressing GFP transfected with mCherry constructs (red) and stained for LAMP1 (grayscale) as well as DAPI (blue) after fixation. Scale bar = 50 µm. **E)** Quantification of GFP distribution based on scheme in Figure 5C. GFP intensity in the different segments did not significantly change upon the expression of the indicated mCherry constructs. Error bars = SEM. **F)** Confocal images of HeLa cells transiently overexpressing GFP or GFP-KIF1B. Cells were immunostained with LAMP1 (grayscale) and PLEKHG3 (red). Scale bar = 20 µm. **G)** GFP immunoblot of the HeLa cells shown in F. Expression of GFP-KIF1B is lower compared to GFP, which might be due to decreased transfer efficiency for high molecular weight proteins during western blotting. Tubulin staining served as a loading control. **H)** Lysosomal distribution in cells overexpressing the constructs used in F. KIF1B overexpression concentrated lysosomes in the periphery, confirming the effect of KIF1A shown in C. Values represent mean of >50 cells from 3 biological replicates. Error bars = SEM. Statistics = Student's t-test $p<0.05$. **I)** Epifluorescence images of HeLa cells transiently overexpressing GFP-KIF1B as shown in F. Cells were immunostained with LAMP1 (grayscale) and PLEKHG3 (red) and imaged on an Olympus SlideScanner for medium throughput analysis. In merged images the nuclei are stained with DAPI and shown in blue. Regions of interest are dotted in red to indicate LAMP1 high, lysosome rich areas, or in green to indicate LAMP1 low, lysosome poor areas. Scale bar = 20 µm. **J-K)** Quantification of experiments similar to the one shown in I. **J)** Superplots represent the average integrated density of PLEKHG3 in the LAMP1 high or low regions (top). The intensity of LAMP1 in these high and low regions is shown at the bottom as a control. **K)** Average integrated density of LAMP1 in the PLEKHG3 high or low regions (top). The intensity of PLEKHG3 in these high and low regions is shown at the bottom as a control. **L-N)** Morphometric analysis of cells shown in Figure 5A. Cells were segmented based on the GFP channel. Morphometric analysis was performed by Fiji's shape descriptor tool. Bars represent three independent experiments. Error bars = SEM. In the superplots in A, J and K the dots represent individual data points of each of three color coded biological replicates; diamonds represent the mean of each replicate; black bars represent the mean ± the SEM of the three biological replicates. In **E and L-N,** black asterisks denote p values according to Student's t-test, comparing the effect of KIF1A or RUFY3 expression with the effect of mCherry. White asterisks in the bars indicate the p values according to Student's t-test comparing GFP-PLEKHG3 and GFP expressing cells.

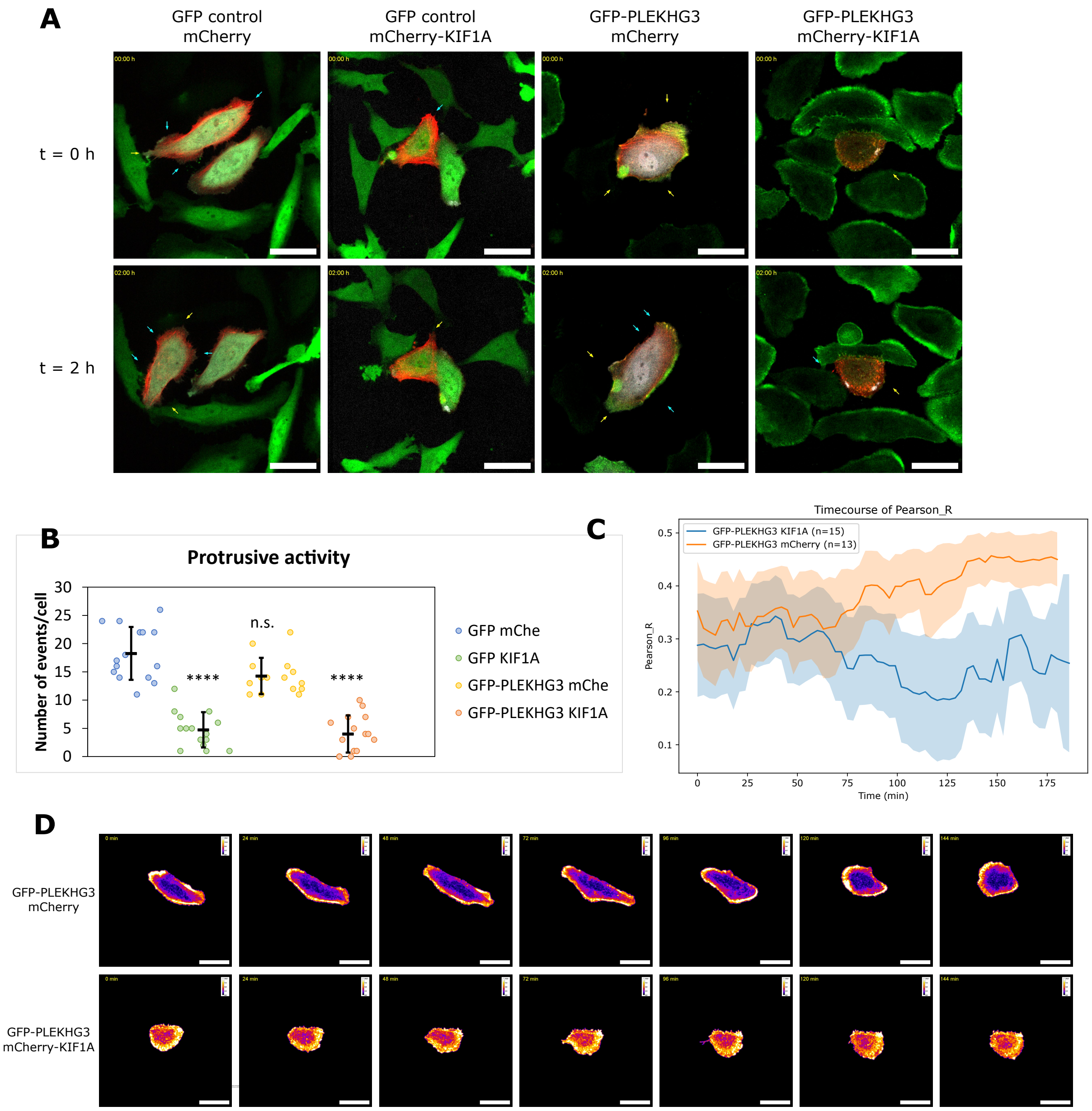
A
GFP control
mCherry
GFP control
mCherry-KIF1A
GFP-PLEKHG3
mCherry
GFP-PLEKHG3
mCherry-KIF1A
t = 0 h
t = 2 h
B
Protrusive activity
Number of events/cell
n.s.
****
****
GFP mChe
GFP KIF1A
GFP-PLEKHG3 mChe
GFP-PLEKHG3 KIF1A
C
Timecourse of Pearson_R
GFP-PLEKHG3 KIF1A (n=15)
GFP-PLEKHG3 mCherry (n=13)
Pearson_R
Time (min)
D
GFP-PLEKHG3
mCherry
GFP-PLEKHG3
mCherry-KIF1A

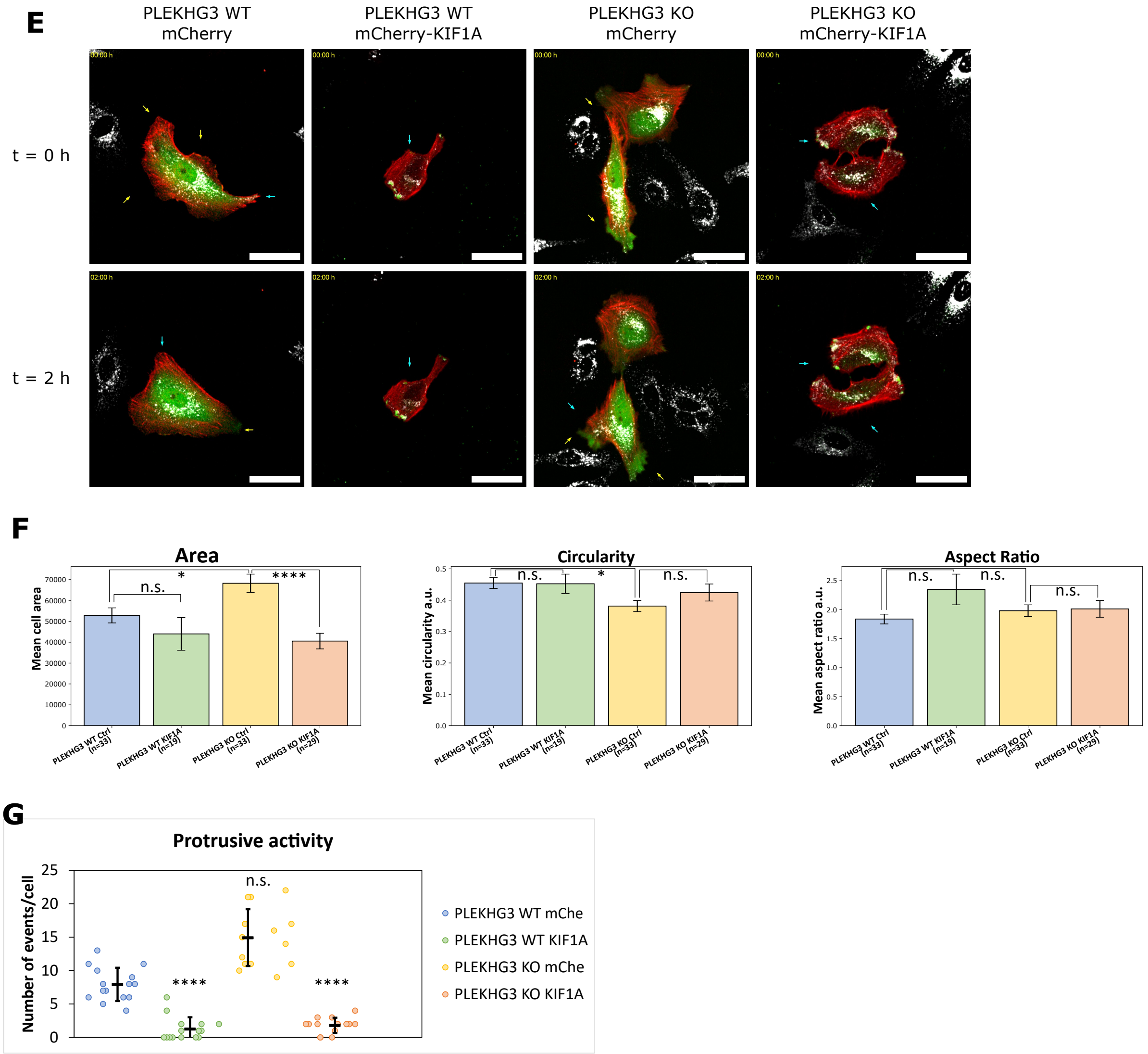


**Figure S6: PLEKHG3 localizes to F-actin independently of lysosomal transport and is dispensable for protrusive activity. A)** Stills from live cell imaging (Movies S12-15). Cells stably expressing GFP or GFP-PLEKHG3 (false color: green) were cotransfected with BFP-LifeAct (false color: red) and the indicated mCherry constructs (false color: grey). Yellow arrows = emerging protrusions; blue arrows = retracting protrusions. Stills were generated over a period of 2 hrs. Scale bar = 50 µm. **B)** Quantification of protrusions formed and retracted over time in cells from A. Values indicate average number of protrusions formed in a timespan of three hours from a total of ≥ 15 cells per condition. Error bars = SEM. **C)** Quantification of GFP-PLEKHG3 and BFP-LifeAct colocalization by Fiji's Coloc2 Plugin (see materials and methods) over a timespan of three hours. Lines represent mean of all cells per condition, and light-color shading represents the SEM. **D)** Stills from heatmap videos from live cell imaging performed in A (Movies S16-17). Heatmaps represent overlap intensity of GFP-PLEKHG3 and LifeAct. **E)** Stills from live cell imaging (Movies S18-21). PLEKHG3 WT and PLEKHG3 KO cells were transfected with the indicated mCherry constructs and incubated with LysoTracker. Yellow arrows = emerging protrusions; blue arrows = retracting protrusions. Stills were generated over a period of 2 hrs. Scale bar = 50 µm. **F)** Morphometric analysis of cells over time. Cell shape analysis was performed with Fiji's shape descriptors (see materials and methods) per cell every three minutes. Bar plots represent the average over all time points to represent changes in shape over time. **G)** Quantification of emerging and retracting protrusions over time in cells from E. Values indicate average number of protrusions in one hour from a total of ≥ 15 cells per condition. Error bars = SEM. In **B,F-G,** black asterisks denote p values according to 1-way-ANOVA or Kruskal-Wallis and Bonferroni post-hoc testing, comparing the effect of KIF1A against mCherry or PLEKHG3 WT against KO.

### Supplementary videos

Movies S1-S4:

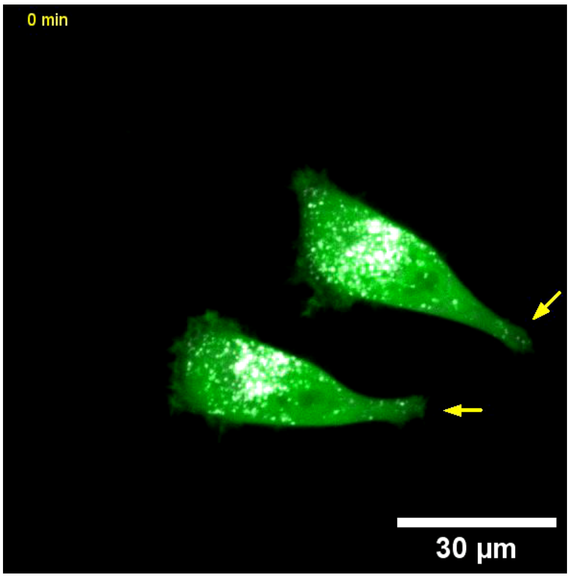


**Movie S1: L3 CTRL HeLa cells stably expressing GFP.** Cells were incubated with LysoTracker Deep Red (False color: grayscale) and imaged for 3h with images taken every 4 min on a live spinning disk microscope (Visitron). Yellow arrows indicate forming and blue arrows retracting protrusions. This video corresponds to stills shown in Figure S3L.

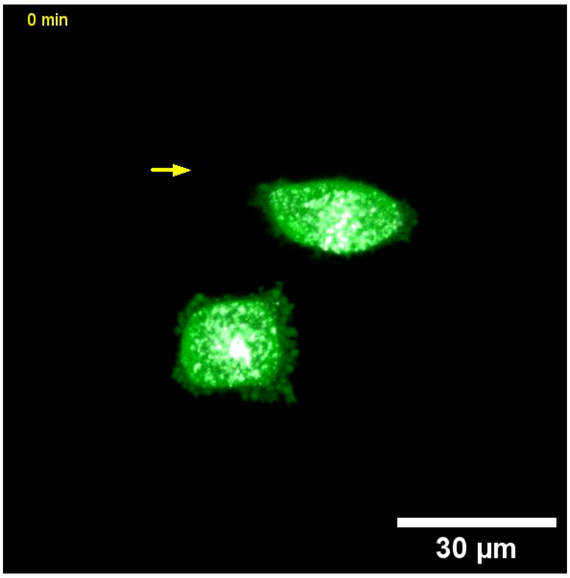


**Movie S2: L3 KO HeLa cells stably expressing GFP.** Cells were incubated with LysoTracker Deep Red (False color: grayscale) and imaged for 3h with images taken every 4 min on a live spinning disk microscope (Visitron). Yellow arrows indicate forming and blue arrows retracting protrusions. This video corresponds to stills shown in Figure S3L

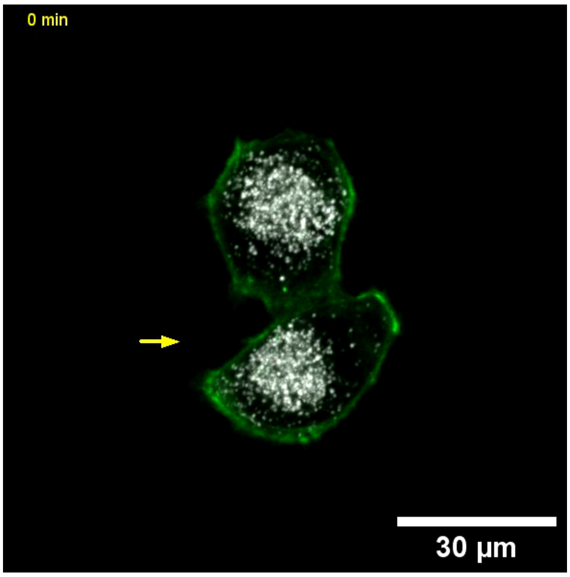


**Movie S3: L3 CTRL HeLa cells stably expressing GFP-PLEKHG3.** Cells were incubated with LysoTracker Deep Red (False color: grayscale) and imaged for 3h with images taken every 4 min on a live spinning disk microscope (Visitron). Yellow arrows indicate forming and blue arrows retracting protrusions. This video corresponds to stills shown in Figure S3L

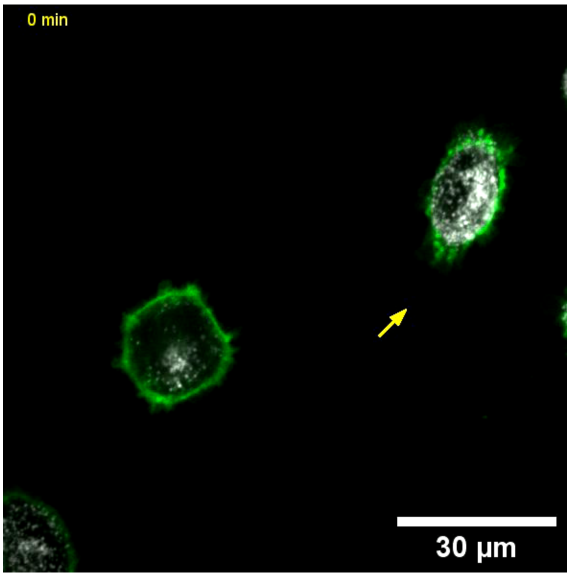


**Movie S4: L3 KO HeLa cells stably expressing GFP-PLEKHG3.** Cells were incubated with LysoTracker Deep Red (False color: grayscale) and imaged for 3h with images taken every 4 min on a live spinning disk microscope (Visitron). Yellow arrows indicate forming and blue arrows retracting protrusions. This video corresponds to stills shown in Figure S3L.

### Movie S5

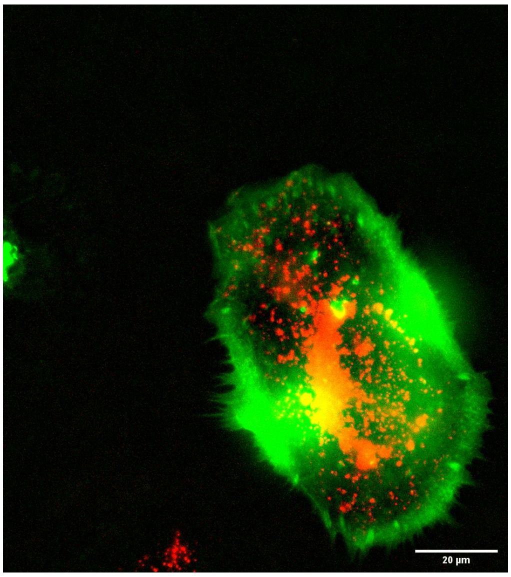


**Movie S5: HeLa cell transfected with GFP-PLEKHG3.** Cells were incubated with LysoTracker Deep Red and imaged for 3h with images taken every 15 min on a Celldiscoverer 7 (Zeiss). Lysosomes concentrate in retracting membrane regions as well as in forming protrusions. This video corresponds to stills shown in Figure 4F.

### Movies S6-11

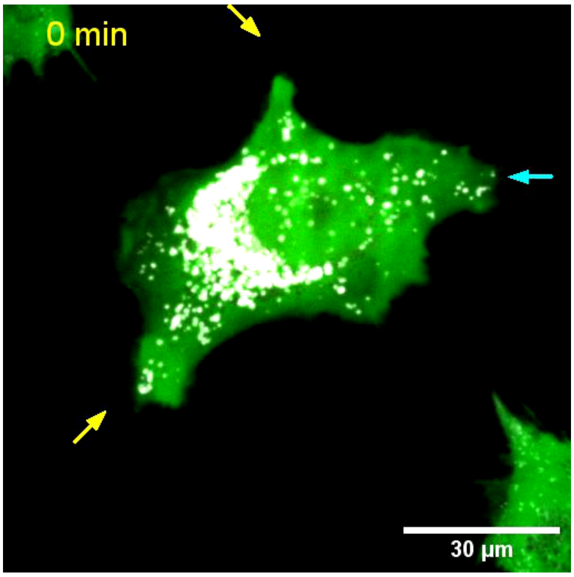


**Movie S6: HeLa cells stably expressing GFP.** Cells were transfected with mCherry (red) and incubated with LysoTracker Deep Red (False color: grayscale) and imaged for 3h with images taken every 4 min on a live spinning disk microscope (Visitron). Yellow arrows indicate forming and blue arrows retracting protrusions. This video corresponds to stills shown in Figure 6A. GFP and LysoTracker channels were acquired in a z-stack while mCherry channel was recorded as a centered slice, which explains the blinking in this channel.

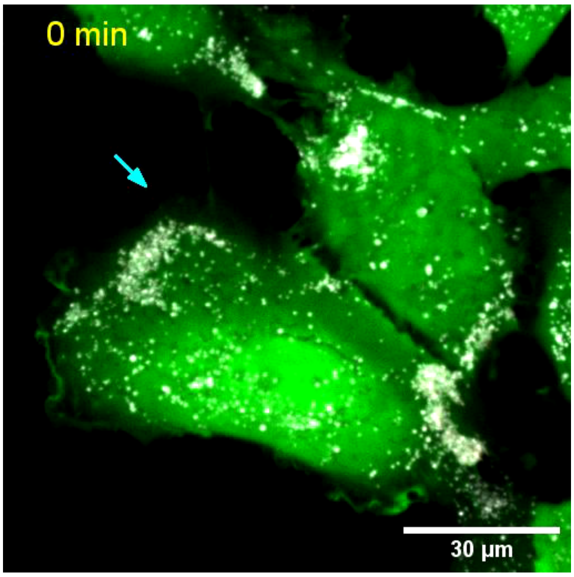


**Movie S7: HeLa cells stably expressing GFP.** Cells were transfected with mCherry-KIF1A (red) and incubated with LysoTracker Deep Red (False color: grayscale) and imaged for 3h with images taken every 4 min on a live spinning disk microscope (Visitron). Yellow arrows indicate forming and blue arrows retracting protrusions. This video corresponds to stills shown in Figure 6A. GFP and LysoTracker channels were acquired in a z-stack while mCherry channel was recorded as a centered slice, which explains the blinking in this channel.

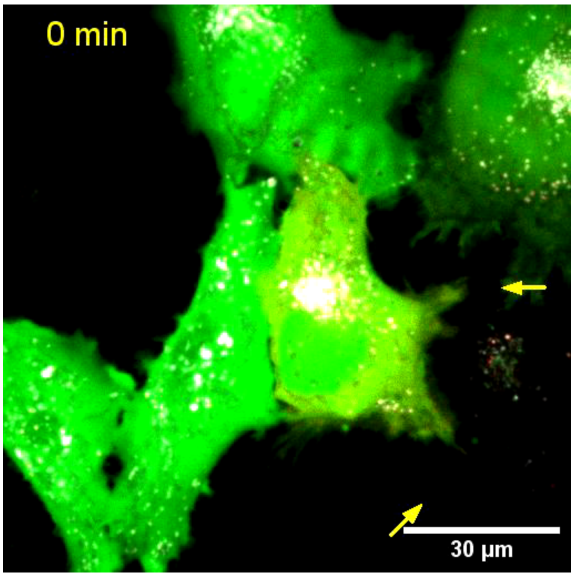


**Movie S8: HeLa cells stably expressing GFP.** Cells were transfected with RUFY3-mCherry (red) and incubated with LysoTracker Deep Red (False color: grayscale) and imaged for 3h with images taken every 4 min on a live spinning disk microscope (Visitron). Yellow arrows indicate forming and blue arrows retracting protrusions. This video corresponds to stills shown in Figure 6A. GFP and LysoTracker channels were acquired in a z-stack while mCherry channel was recorded as a centered slice, which explains the blinking in this channel.

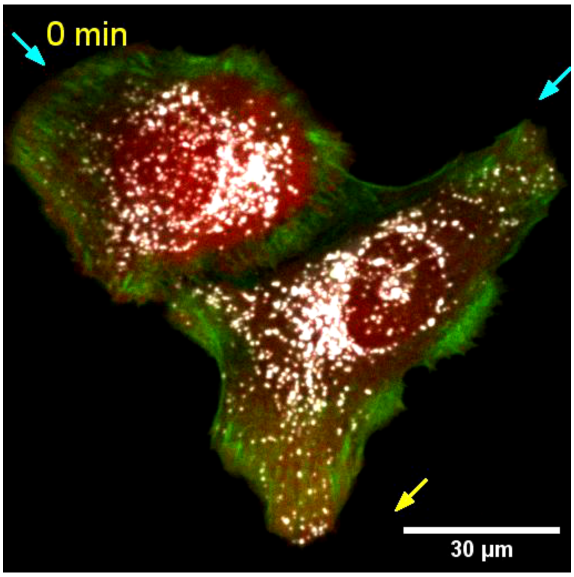


**Movie S9: HeLa cells stably expressing GFP-PLEKHG3.** Cells were transfected with mCherry (red) and incubated with LysoTracker Deep Red (False color: grayscale) and imaged for 3h with images taken every 4 min on a live spinning disk microscope (Visitron). Yellow arrows indicate forming and blue arrows retracting protrusions. This video corresponds to stills shown in Figure 6A. GFP and LysoTracker channels were acquired in a z-stack while mCherry channel was recorded as a centered slice, which explains the blinking in this channel.

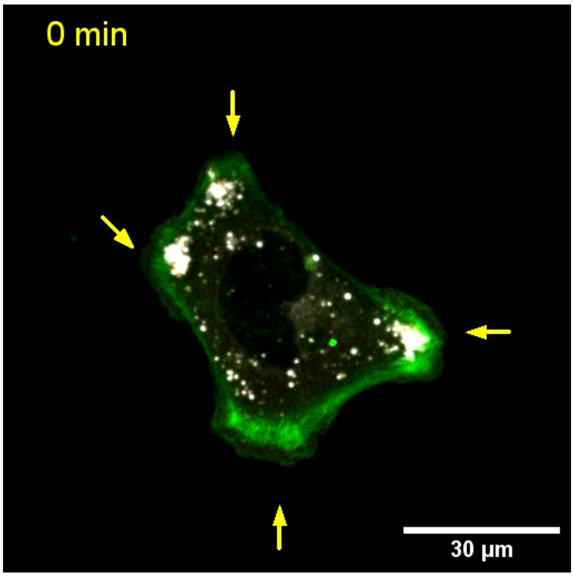


**Movie S10: HeLa cells stably expressing GFP-PLEKHG3.** Cells were transfected with mCherry-KIF1A (red) and incubated with LysoTracker Deep Red (False color: grayscale) and imaged for 3h with images taken every 4 min on a live spinning disk microscope (Visitron). Yellow arrows indicate forming and blue arrows retracting protrusions. This video corresponds to stills shown in Figure 6A. GFP and LysoTracker channels were acquired in a z-stack while mCherry channel was recorded as a centered slice, which explains the blinking in this channel.

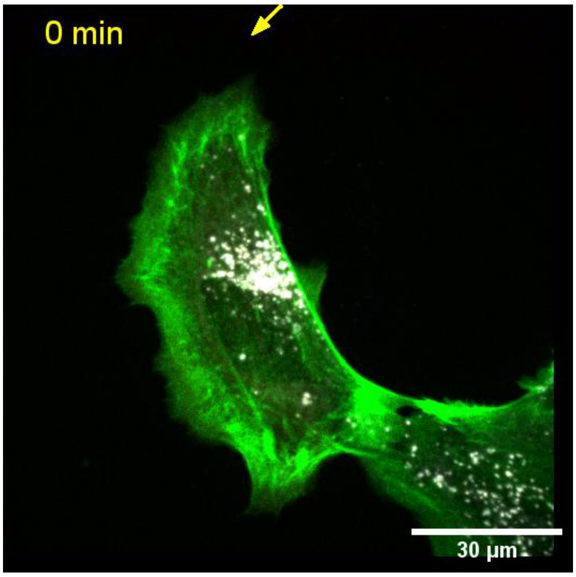


**Movie S11: HeLa cells stably expressing GFP-PLEKHG3.** Cells were transfected with RUFY3-mCherry (red) and incubated with LysoTracker Deep Red (False color: grayscale) and imaged for 3h with images taken every 4 min on a live spinning disk microscope (Visitron). Yellow arrows indicate forming and blue arrows retracting protrusions. This video corresponds to stills shown in Figure 6A. GFP and LysoTracker channels were acquired in a z-stack while mCherry channel was recorded as a centered slice, which explains the blinking in this channel.

### Movies S12-15

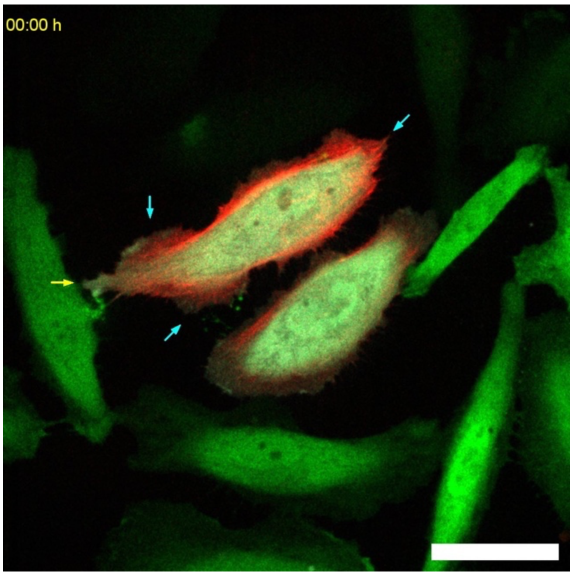


**Movie S12: HeLa cells stably expressing GFP.** Cells were transfected with mCherry (False color: grayscale) and BFP-LifeAct (False color: red) and imaged for 3h with images taken every 3 min on a live spinning disk microscope (Visitron). Yellow arrows indicate forming and blue arrows retracting protrusions. This video corresponds to stills shown in Figure S6A. All channels were acquired in a z-stack and maximum projected for better visualization.

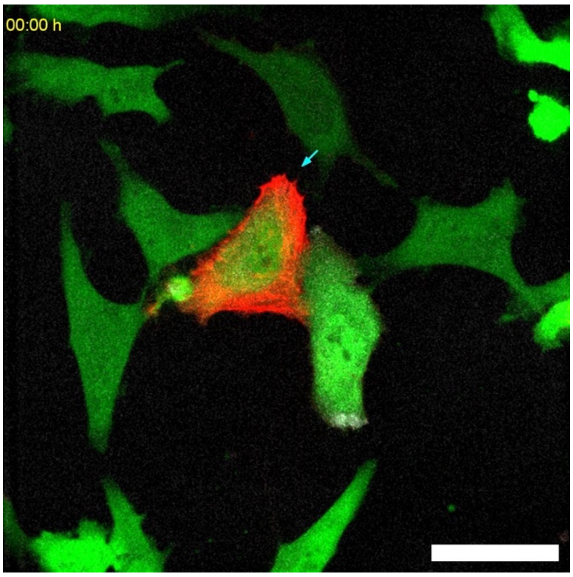


**Movie S13: HeLa cells stably expressing GFP.** Cells were transfected with mCherry-KIF1A (False color: grayscale) and BFP-LifeAct (False color: red) and imaged for 3h with images taken every 3 min on a live spinning disk microscope (Visitron). Yellow arrows indicate forming and blue arrows retracting protrusions. This video corresponds to stills shown in Figure S6A. All channels were acquired in a z-stack and maximum projected for better visualization.

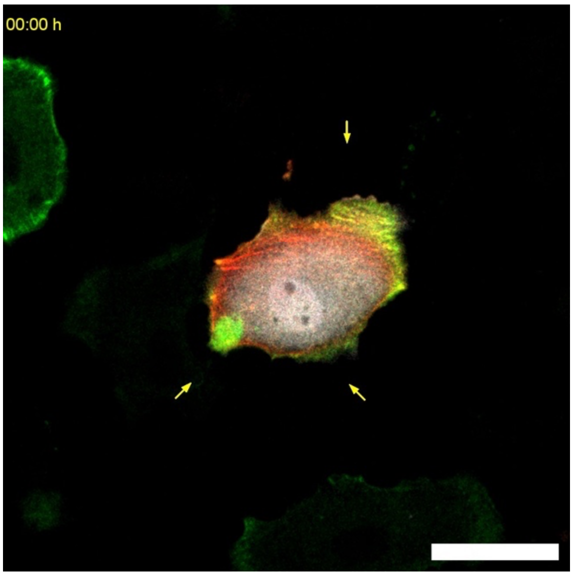


**Movie S14: HeLa cells stably expressing GFP-PLEKHG3.** Cells were transfected with mCherry (False color: grayscale) and BFP-LifeAct (False color: red) and imaged for 3h with images taken every 3 min on a live spinning disk microscope (Visitron). Yellow arrows indicate forming and blue arrows retracting protrusions. This video corresponds to stills shown in Figure S6A. All channels were acquired in a z-stack and maximum projected for better visualization.

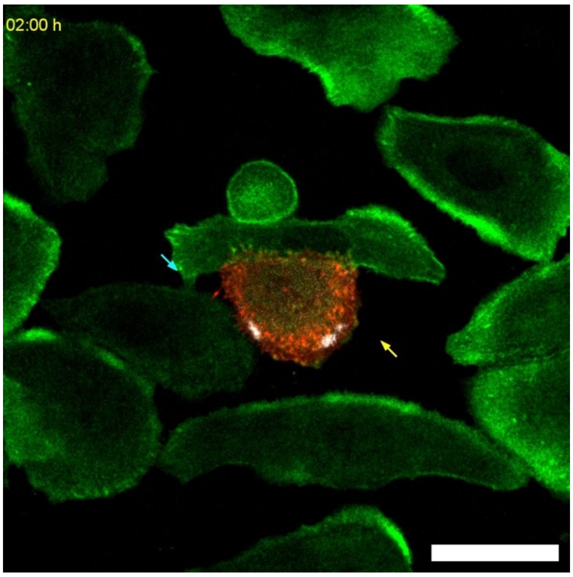


**Movie S15: HeLa cells stably expressing GFP-PLEKHG3.** Cells were transfected with mCherry-KIF1A (False color: grayscale) and BFP-LifeAct (False color: red) and imaged for 3h with images taken every 3 min on a live spinning disk microscope (Visitron). Yellow arrows indicate forming and blue arrows retracting protrusions. This video corresponds to stills shown in Figure S6A. All channels were acquired in a z-stack and maximum projected for better visualization.

### Movies S16-17

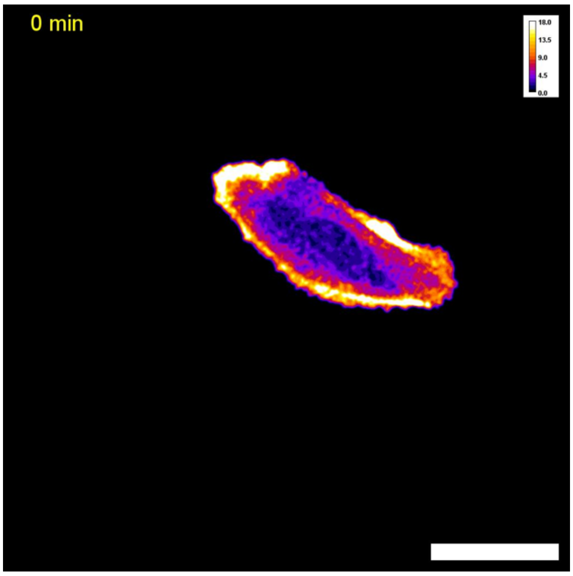


**Movie S16: Heatmap video of HeLa cell expressing GFP-PLEKHG3 and LifeAct.** Cells are taken from the experiment in Movies S 12-15. Cells were transfected with mCherry control and imaged for 3h with images taken every 3 min on a live spinning disk microscope (Visitron). The cell was segmented and the background around the cell's outline cleared. The heatmap represents the colocalization intensity of GFP-PLEKHG3 and BFP-LifeAct. This video corresponds to stills shown in Figure S6D.

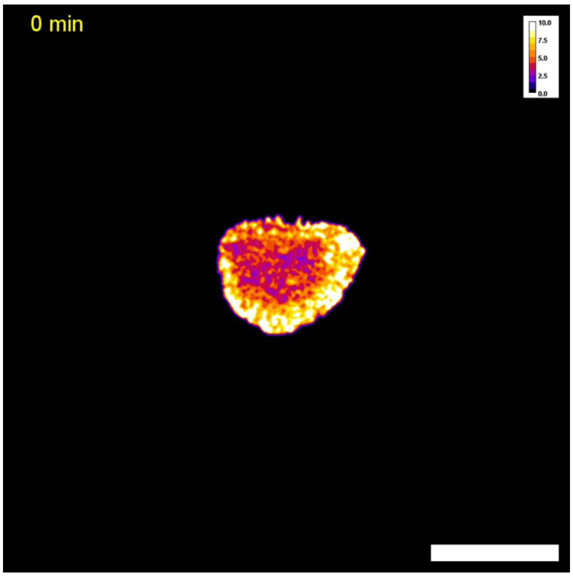


**Movie S17: Heatmap video of HeLa cell expressing GFP-PLEKHG3 and LifeAct.** Cells are taken from the experiment in Movies S 12-15 (same cell as in Movie S15). Cells were transfected with mCherry-KIF1A and imaged for 3h with images taken every 3 min on a live spinning disk microscope (Visitron). The cell was segmented and the background around the cell's outline cleared. The heatmap represents the colocalization intensity of GFP-PLEKHG3 and BFP-LifeAct. This video corresponds to stills shown in Figure S6D.

### Movies S18-21

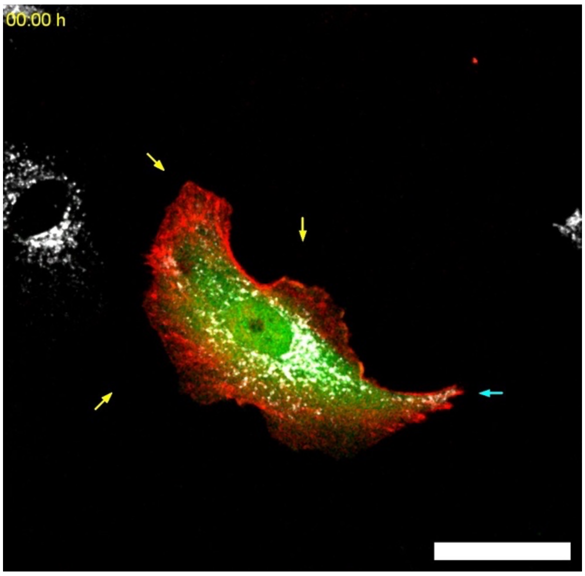


**Movie S18: HeLa PLEKHG3 WT cells.** Cells were transfected with mCherry (False color: green) and BFP-LifeAct (False color: red), incubated with LysoTracker Green (False color: grayscale) and imaged for 3h with images taken every 3 min on a live spinning disk microscope (Visitron). Yellow arrows indicate forming and blue arrows retracting protrusions. This video corresponds to stills shown in Figure S6E. All channels were acquired in a z-stack and maximum projected for better visualization.

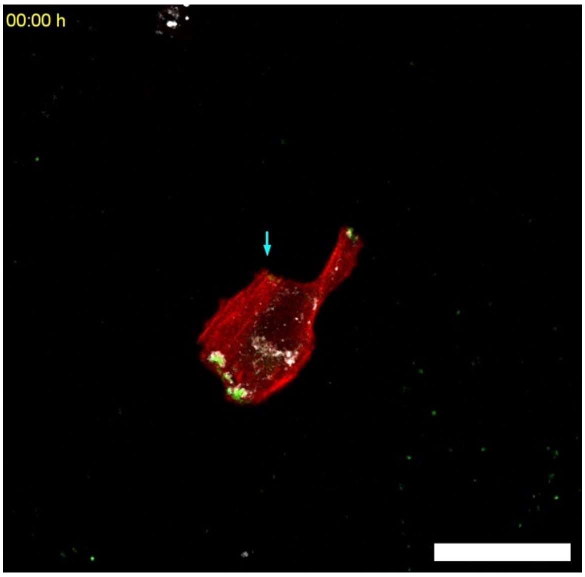


**Movie S19: HeLa PLEKHG3 WT cells.** Cells were transfected with mCherry-KIF1A (False color: green) and BFP-LifeAct (False color: red), incubated with LysoTracker Green (False color: grayscale) and imaged for 3h with images taken every 3 min on a live spinning disk microscope (Visitron). Yellow arrows indicate forming and blue arrows retracting protrusions. This video corresponds to stills shown in Figure S6E. All channels were acquired in a z-stack and maximum projected for better visualization.

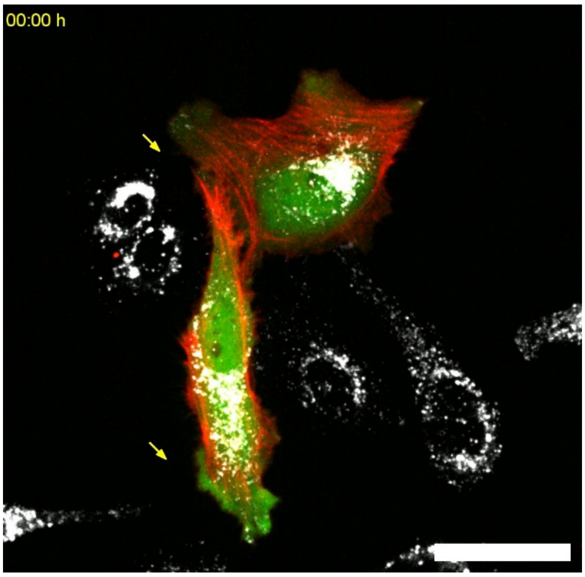


**Movie S20: HeLa PLEKHG3 KO cells.** Cells were transfected with mCherry (False color: green) and BFP-LifeAct (False color: red), incubated with LysoTracker Green (False color: grayscale) and imaged for 3h with images taken every 3 min on a live spinning disk microscope (Visitron). Yellow arrows indicate forming and blue arrows retracting protrusions. This video corresponds to stills shown in Figure S6E. All channels were acquired in a z-stack and maximum projected for better visualization.

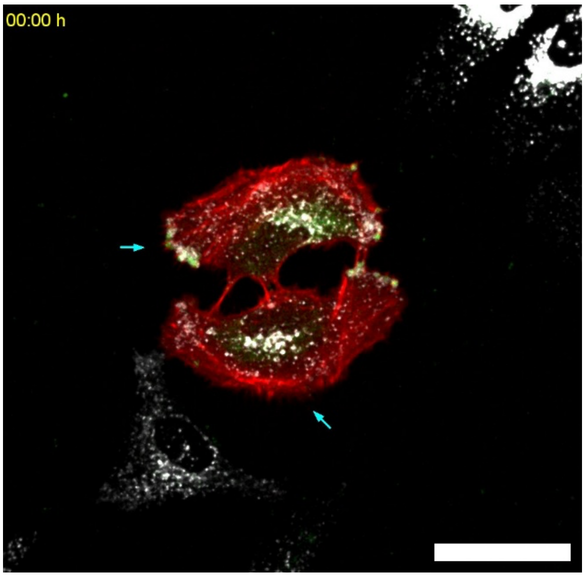


**Movie S21: HeLa PLEKHG3 KO cells.** Cells were transfected with mCherry-KIF1A (False color: green) and BFP-LifeAct (False color: red), incubated with LysoTracker Green (False color: grayscale) and imaged for 3h with images taken every 3 min on a live spinning disk microscope (Visitron). Yellow arrows indicate forming and blue arrows retracting protrusions. This video corresponds to stills shown in Figure S6E. All channels were acquired in a z-stack and maximum projected for better visualization.

### Supplementary Tables

**Table S1: All identified proteins and list of proteins according to Figure S1D.** Contains all identified proteins, spectral counts, LFQ intensities, and differential expression analysis results for L3-T-V5, i.e. hits from L3-T-V5 bait significantly enriched over GFP-V5-CYTO (log2FC ≥ 1.49 and padj ≤ 0.05). Nuclear hits were removed by excluding preys with GO-term 'nucleus' from the list if not enriched in at least one condition of the V5-LYSO bait over the V5-CYTO control (log2FC ≥ 1.49 and padj ≤ 0.05) from second screen.

Supplementary Table 1_L3-T-V5.xlsx

**Table S2: All identified proteins and list of proteins according to Figure S1I-K.** Contains all identified proteins, spectral counts, LFQ intensities, and differential expression analysis results for V5-T-L3, i.e. hits from V5-T-L3 bait significantly enriched over V5-CYTO (log2FC ≥ 1.49 and padj ≤ 0.05). Nuclear hits were removed by excluding preys with GO-term 'nucleus' from the list if not enriched in at least one condition of the V5-LYSO bait over the V5-CYTO control (log2FC ≥ 1.49 and padj ≤ 0.05) from this screen.

Supplementary Table 2_V5-T-L3.xlsx
