## Supplementary figures and images for "Peripheral lysosomes recruit PLEKHG3 to focal adhesions and restrain protrusion dynamics"

### Graphical abstract

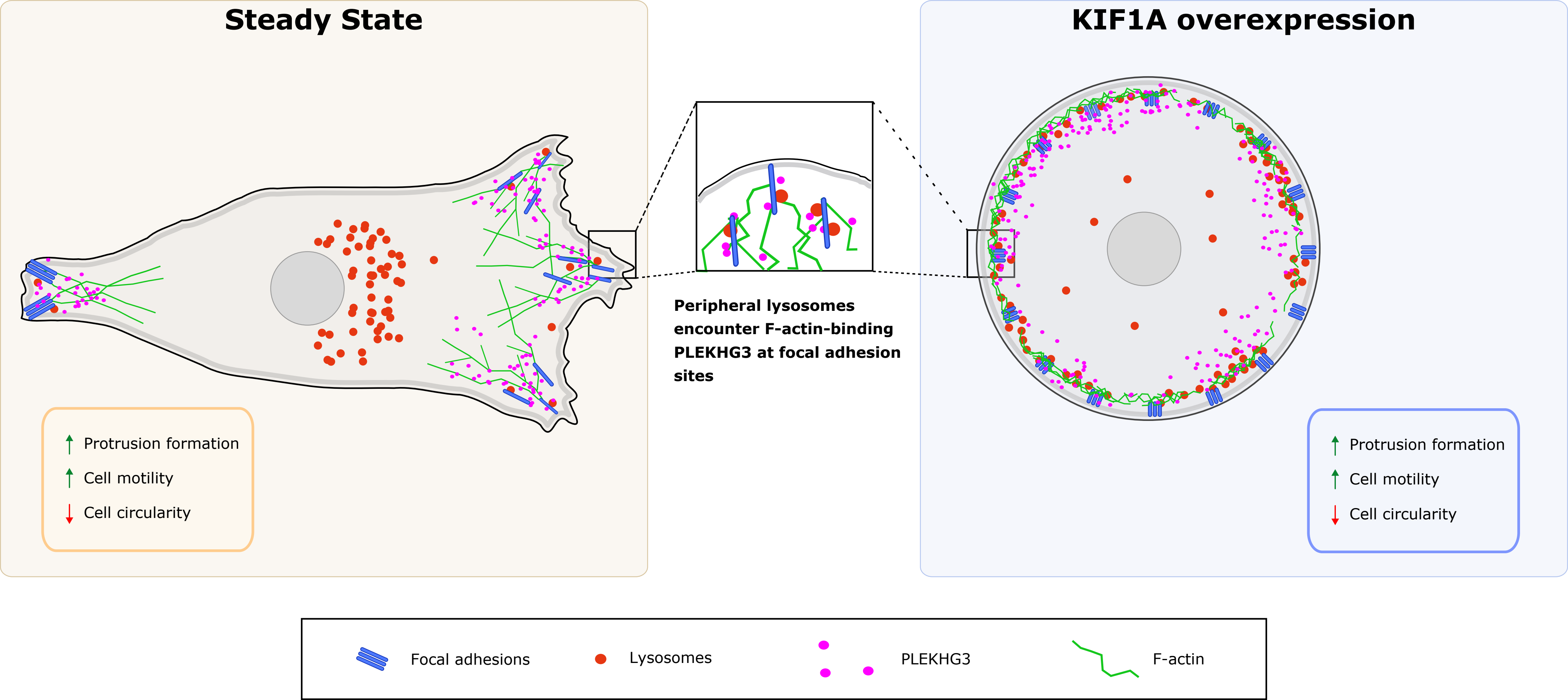
